# Grand Biological Universe: Genome space geometry unravels looking for a single metric is likely to be futile in evolution

**DOI:** 10.1101/2023.07.08.548189

**Authors:** Nan Sun, Hongyu Yu, Ruohan Ren, Tao Zhou, Mengcen Guan, Leqi Zhao, Stephen S.-T. Yau

**Affiliations:** Department of Mathematical Sciences, Tsinghua University, Beijing, China; Zhili College, Tsinghua University, Beijing, China; Yanqi Lake Beijing Institute of Mathematical Sciences and Applications, Beijing, China

**Keywords:** Genome Space, Natural Metric, Geometry, Seven Kingdoms, Grand Biological Universe

## Abstract

Understanding the differences between genomic sequences of different lives is crucial for biological classification and phylogeny. Here, we downloaded all the reliable sequences of the seven kingdoms and determined the dimensions of the genome space embedded in the Euclidean space, along with the corresponding Natural Metrics. The concept of the Grand Biological Universe is further proposed. In the grand universe, the convex hulls formed by the universes of seven kingdoms are mutually disjoint, and the convex hulls formed by different biological groups within each kingdom are mutually disjoint. This study provides a novel geometric perspective for studying molecular biology and also offers an accurate way for large-scale sequence comparison in a real-time manner. Most importantly, this study shows that, due to the space-time distortion in the biological genome space similar to Einstein’s theory, it is futile to look for a single metric to measure different biological universes, as previous studies have done.

## 1. Introduction

Imitating Hilbert who proposed 23 problems in mathematics in 1900, Defense Advanced Research Projects Agency (DARPA) proposed 23 problems in pure and applied mathematics in 2008 [1]. These problems will be proven to be very influential for the development of mathematics in the 21st century. In the DARPA problems, we are asked to understand “The Geometry of Genome Space” (number 15) and “What are the Fundamental Laws of Biology” (number 23). Our convex hull principle for molecular biology states that the convex hull formed from Natural Vectors of one biological group does not intersect with the convex hull formed from any other biological group [2] [3]. This can be viewed as one of the Fundamental Laws of Biology for which DARPA has been looking for since 2008. On the basis of the convex hull principle, we can construct the geometry of the genome space. A genome space consists of all known genomes of living beings and provides insights into their relationships [2]. The genome space can be considered as the moduli space in mathematics, and genome sequences can be canonically embedded in a high-dimensional Euclidean space by means of Natural Vectors [4]. In this space, a sequence is uniquely represented as a point by means of the nucleotide distribution information within the sequence. Similar sequences lie closely, and convex hulls of different groups are disjoint according to the convex hull principle. The geometry of space is reflected in the similarity of sequences. The concept of the genome space is reminiscent of the vast universe and galaxies that exist above us, where different DNA sequences are akin to stars. Chromosomes belonging to the same family form galaxies, and the distances among stars can be found through suitable measurements. The similarity of sequences can be measured by the Natural Metric, which is different from the induced metric from the ambient Euclidean space. In the realm of physics, Einstein’s theory postulates that stars possess the capacity to distort space, thereby engendering distinct metrics within different regions. Concurrently, the pervasive existence of imperceptible dark matter and dark energy contributes further to this spatial distortion. Analogously, the genome space manifests a comparable phenomenon. Within each kingdom, a large number of undetected sequences coalesce with known sequences, profoundly influencing the underlying spatial structure and, subsequently, yielding distinct geometries for each kingdom. Our study substantiates this notion, as we reveal substantial variations in metrics across each genome space, underscoring the parallelism between the physical and genetic realms.

With the progress of sequencing technology, more and more sequences have been sequenced. According to the records of NCBI (National Center for Biotechnology Information, https://www.ncbi.nlm.nih.gov/genbank/statistics/), the number of sequences has increased exponentially.

These sequences include complete chromosomes or organelle, gene fragments, or protein sequences. A genome is the sum of all genetic material of an individual [5]. A genome contains genes of chromosomes and organelles, etc, and chromosomes carry part or all genetic information [6]. Almost all eukaryotic species contain nuclear genome (chromosome) and organelle genome (mitochondria), plants also have chloroplast DNA. Most Eukaryotes have multiple linear chromosomes. Prokaryotes include archaea and bacteria, and their genetic material is mainly carried by chromosomes. For some bacteria, the auxiliary genetic material in the plasmid is also part of the genome [7]. Archaea and most bacteria genomes have a single circular chromosome [8], but some bacteria have linear or multiple chromosomes [9][10]. The genome size and the chromosome number of eukaryotes and prokaryotes have enormous differences. Escherichia coli O157:H7 strain Sakai has 1 chromosome and 2 plasmids, and its genome size is about 5.6 Mbp (GCF_000008865.2 Strain Sakai substr. RIMD 0509952). Haploid yeast has 16 chromosomes, and its genome size is about 12 Mbp (GCA_000146045.2 R64). Hexaploid bread wheat has 42 chromosomes, and its genome size is about 14454 Mbp (GCA_018294505.1 IWGSC CS RefSeq v2.1). Diploid human has 46 chromosomes, of which 24 linear molecules comprise 3.1 billion nucleotides (GCF_000001405.40 GRCh 38.p14). Invertebrates have small genomes, fish and amphibians have intermediate-size genomes, and birds have relatively small genomes. An important problem is how to compare these diverse sequences.

Sequence alignment is one of the main tools for upstream analysis of biomolecular sequencing data, and there are alignment-based and alignment-free sequence analysis approaches [11]. The pioneering approaches for sequence analysis were based on sequence alignment either global or local, pairwise or multiple sequence alignment [12][13]. Alignment-based approaches generally give excellent results when the sequences under study are closely related and can be reliably aligned, but a reliable alignment cannot be obtained when sequences are much divergent. Another drawback of the alignment is that the addition of new sequences necessitates the process of realignment in multiple sequence alignment. Further limitations are their computational complexity and time-consuming. These alignment-based algorithms are not efficient enough and unable to obtain similarity information between large-scale sequences rapidly. To overcome these limitations, alignment-free methods are developed, and they are mainly based on k-mer frequency, the length of common substrings, the number of word matches, micro-alignments, information theory and graphical representation [11][14]. In recent years, our team has developed various alignment-free algorithms, such as improved k-mer Natural Vector [2], gene graphical representation [15], Fourier transform frequency spectrum [16], and other machine learning based method [17]. These methods provide a very efficient and accurate way to compare large biological datasets. Important practical applications include human origin [18], COVID-19 traceability [19], and protein sequence identification [20]. Excellent theoretical applications include definitions of virus genome space [2], protein space [21], and natural distance [22], which reveal the geometric structure of biological sequences. Specifically, our algorithm (For example, k-mer Natural Vector) can be used to describe genomic sequence data as numerical vectors, and then these numerical vectors can be compared in a geometric space. Similar sequences have similar vector distributions. In this way, like galaxies in the universe, each genome sequence has a clear location and each biological group lies closely in a proper mathematical space. This provides an innovative idea for solving problems 15 and 23 in the list of 23 mathematical challenges. It should be noted that some methods use relative distances (the distance between sample sequence and reference sequences) as vectors to construct a space [23], but this is not a genome space, because the sequence and the point are not in one-to-one correspondence. In addition, except for the genome space for evolution, the three-dimensional spatial structure of the genome (especially eukaryotic chromatin) is also of great significance (but should be distinguished with genome space). Some studies hope to improve its resolution [24], but this is not what we want to focus on.

In our project, we download all reliable chromosome sequences of seven kingdoms (bacteria, archaea, fungi, plant, protozoa, vertebrate, invertebrate) in NCBI to construct the geometric structure of genome space and validate the convex hull principle of genomes. For each kingdom, sequences can be characterized by k-mer Natural Vectors, and each biological group can form a convex hull through k-mer Natural Vectors.

By verifying the convex hull principle, we adjust k and the order of central moment of k-mer Natural Vector, and embed the genome space into the high-dimensional Euclidean space. The weighted sum of Euclidean distance of k-mer Natural Vector is utilized to describe the similarity of sequence structure and function, the weight is determined by the nearest neighborhood classification accuracy. In order to integrate all the sequences of the seven kingdoms into the same space, the concept of the Grand Biological Universe is conceived. The dimension of the Grand Biological Universe is the maximum of all universe dimensions, and the metric can measure the distance of each two universes. To sum up, genome space with a proper metric is a powerful means to determine the phylogeny and classification of genomes. This research has made a key breakthrough in bioinformatics and provided a new geometric perspective for biology.

## 2. Results

### 2.1. The dimension of each kind biological genome space embedded in Euclidean space

Understanding biological classification is necessary for our task, as shown in Figure 1. We use sequence datasets of the seven kingdoms to validate our genome universe theory. (Details about the seven kingdoms are shown in Materials and Methods.) We first construct the genome space for each kingdom, the pipeline is illustrated in Figure 2. The example is based on 3 families and k=1. To begin with, we transform each sequence of the three families into a one-to-one corresponding (4+4n)-dimensional natural vector with high order central moment (Suppose n is a constant and n ∈ Z^+^). Then 3 convex hulls are formed. The genome space is constructed based on the convex hull principle of genomes, which points out that convex hulls corresponding to different biological groups do not overlap with each other. If the 3 convex hulls are disjoint mutually in (4+4n)-dimensional Euclidean space, the genome space exists and can be embedded in (4+4n)-dimensional Euclidean space. Otherwise, n is adjusted until the convex hull principle holds in a certain space. In this space, each point represents a Natural Vector of a genome, and each biological group corresponds to a point cloud, but how these sequences are arranged is unknown. The convex hull principle explains the arrangement of these sequences in the genome space: sequences with similar nucleotide distributions lie in the same convex hull, convex hulls corresponding to different families are disjoint, and all convex hulls show the global landscape of genomes at the family or other classification level.

**Figure 1:**
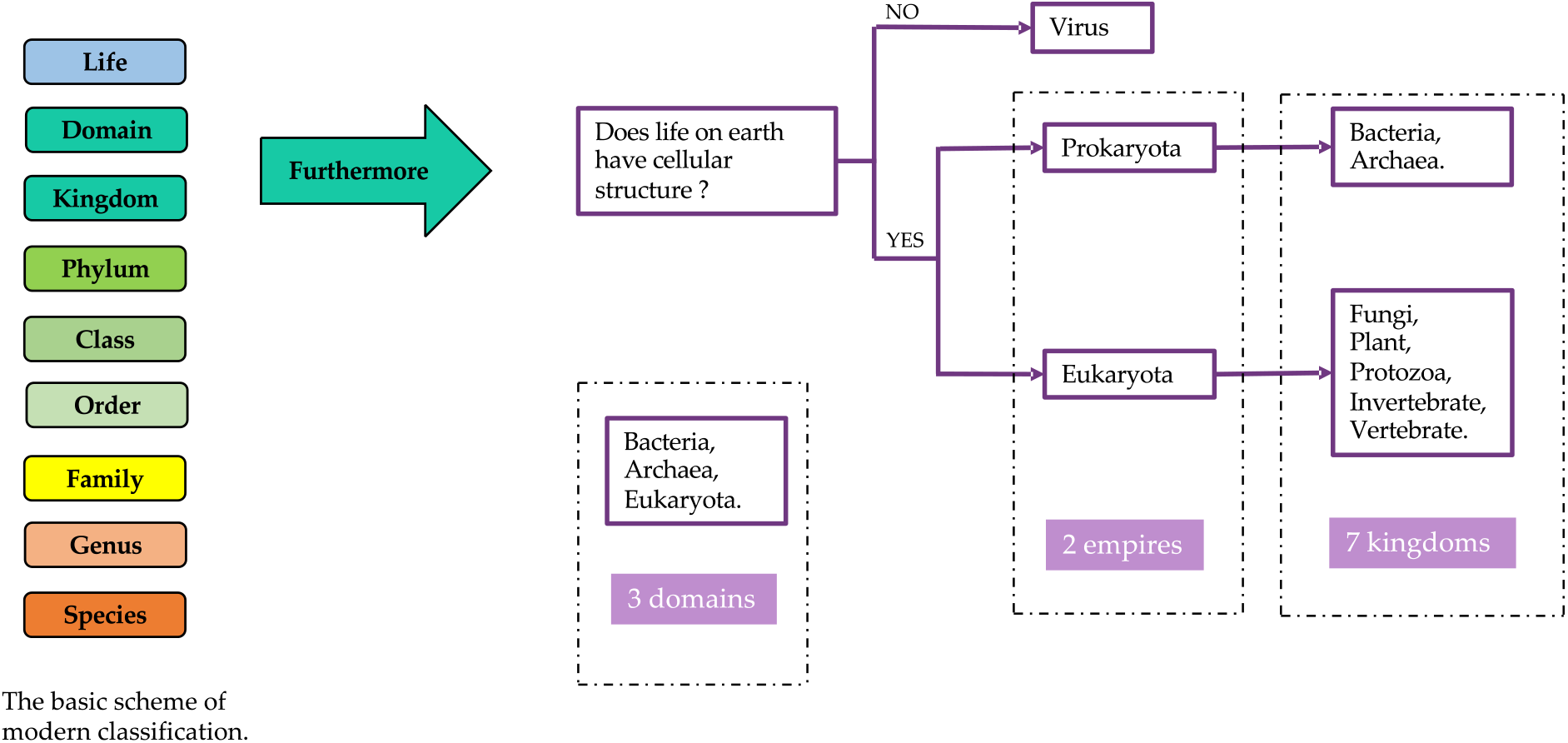
Modern classification. The principal ranks of modern classification and the classification label according to whether life on earth has cellular structure.

**Figure 2:**
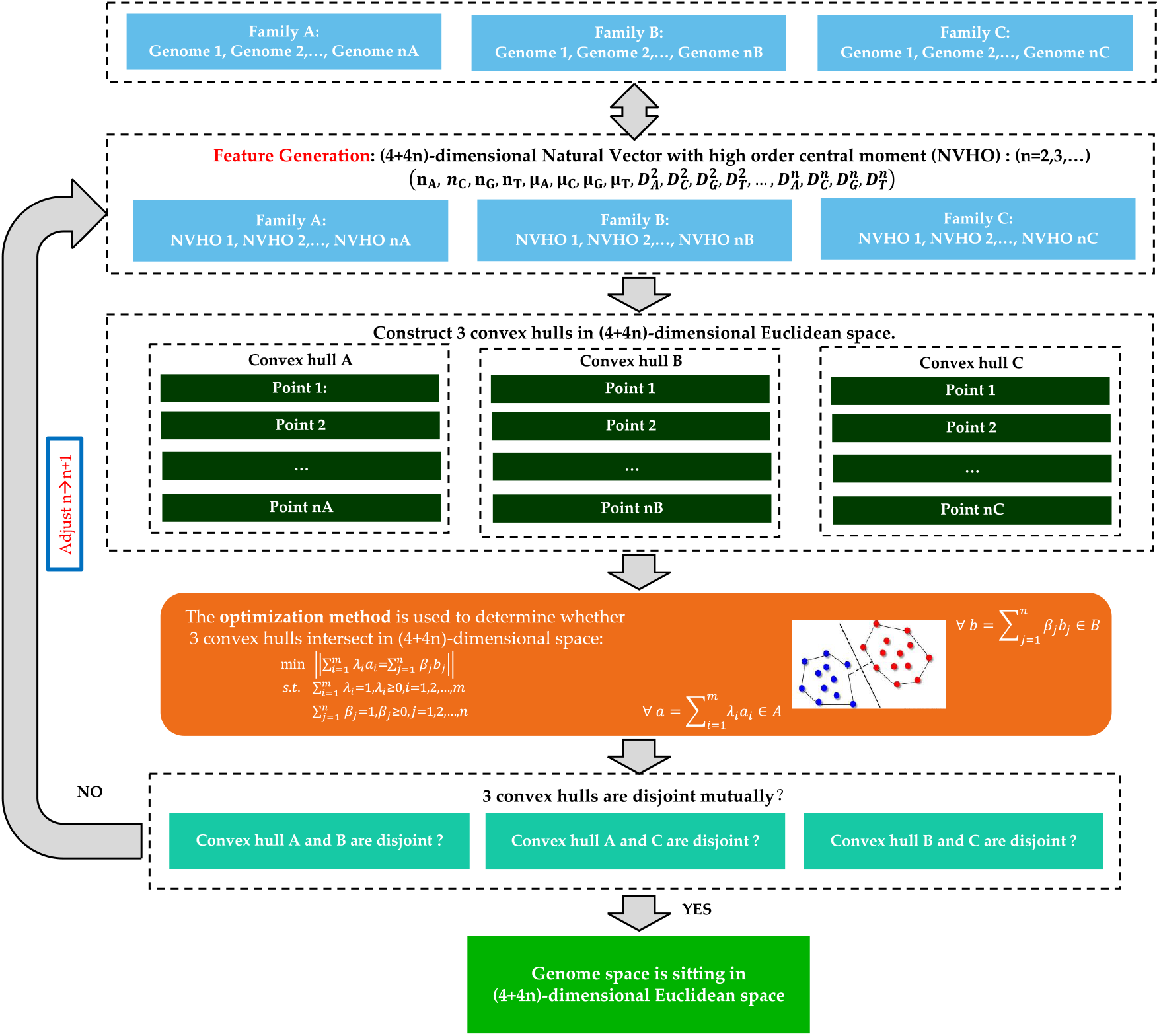
Flowchart of constructing the genome space. The example is based on 3 families and k=1. All convex hulls are mutually disjoint in a (4+4n)-dimensional space (n=2, 3, 4…).

Column B of Table 1 exhibits the dimension of Euclidean space in which the convex hull principle of genomes of each kingdom holds. The largest dimension of all universes is 48, which corresponds to the NV with 11-order moment of bacteria and 2-mer NV with 2-order moment of archaea and invertebrates. The smallest dimension of all universes is 24, which corresponds to the NV with 5-order moment of the plants. The dimension of each biological genome space in Column B represents the corresponding dimension embedded in Euclidean space, which is determined according to the definition of embedding dimension of the moduli space. We test the convex hull principle in spaces of different dimensions, and choose the space with the lowest dimension as the dimension of genome space. The calculation details can be found in Tables A3.1 to A3.7 of Appendix 3.

**Table 1:**
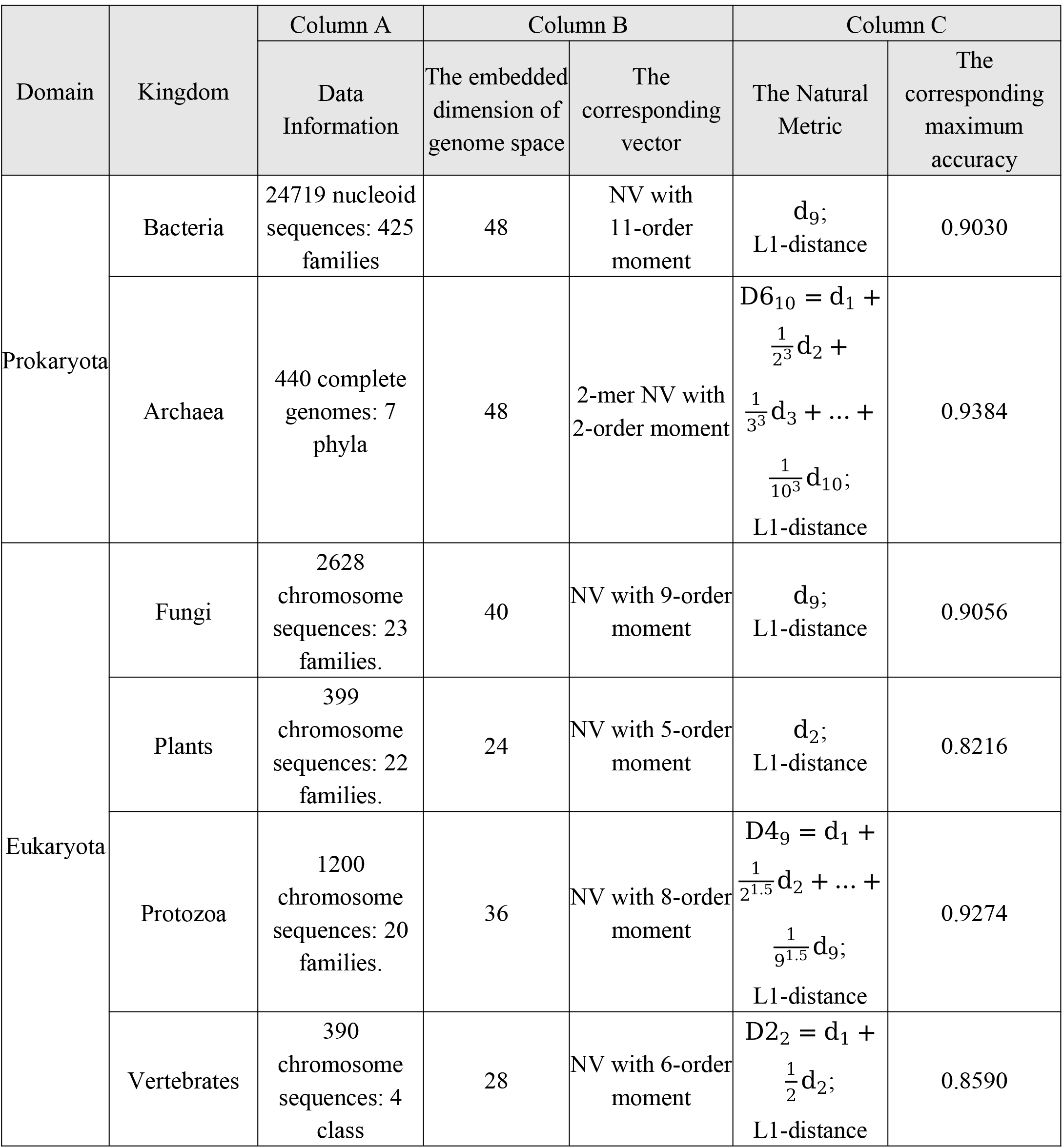
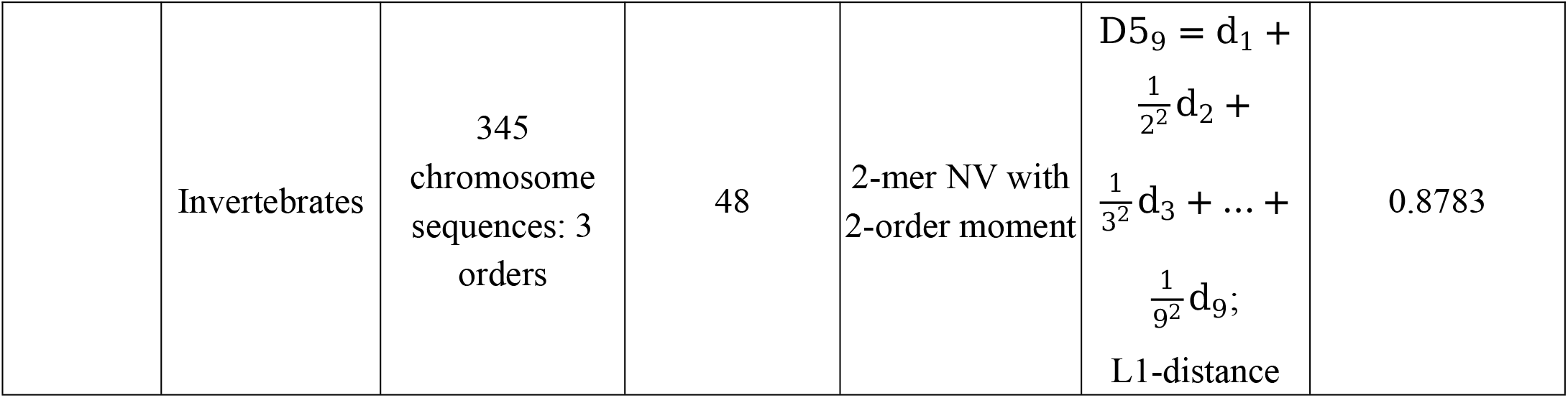
Geometric construction of genome space of seven kingdoms based on chromosomal sequences. Seven kingdoms include Bacteria, Archaea, Fungi, Plants, Protozoa, Vertebrates, and Invertebrates. Column A: Summary information of seven biological sequence datasets. Detailed information can be found in Data A2.1 to A2.7 of Appendix 2. Column B: The dimension of each kind biological genome space embedded in Euclidean space. The calculation details can be found in Tables A3.1 to A3.7 of Appendix 3. Column C: The Natural Metric determination in each kind biological genome space. The calculation detail can be found in Tables A4.1 to A4.7 of Appendix 4.

### 2.2. The Natural Metric determination in each kingdom’s biological genome space

To show the geometry of the genome space of each kingdom, a suitable and descriptive metric is needed. The metric is expected to reflect the dissimilarity among sequences in a good way, which has important implications for geometric biology. Here we suppose the mathematical distance of the corresponding k-mer Natural Vectors (x and y) can well measure the biological distance between two sequences. The institutive metric is Euclidean distance of k-mer Natural Vectors (L1-distance 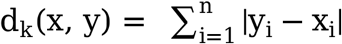, L2-distance 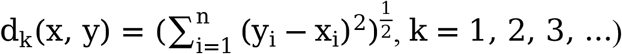, which has been commonly used in previous studies [2][3][4]. Further consideration is the weighted summation of Euclidean distance of k-mer Natural Vectors: D_n_(x, y) = a_1_d_1_(x, y) + a_2_d_2_(x, y) + a_3_d_3_(x, y) + … + a_n_d_n_ x, y, a_k_ ∈ R^+^, k = 1, 2, …, n. D_n_ creatively contains the distribution information from 1-mer to n-mer. a_k_ reflects the degree of contribution of k-mer NV to the 1NN accuracy. The uncertainty of k and weight a_k_ provides space for improving classification accuracy. Normally, we can test the following a_k_:

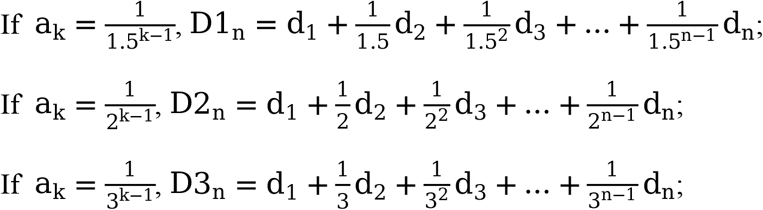

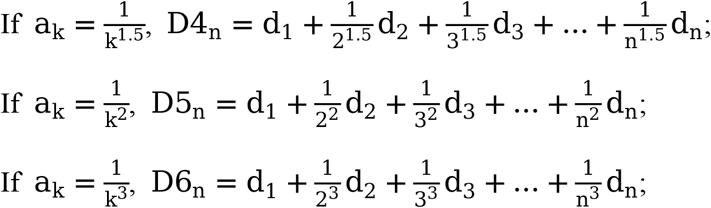

The steps of determining appropriate Natural Metrics are shown in Figure 3. The example is based on 3 families and 12 sequences. Each sequence is converted into a k-mer NV first, and the mutual distance can be obtained through the geometric metric. Then 1NN classification analysis is performed using the distance matrix. For each sequence i, the nearest sequence j can be determined. If sequence i and sequence j share the same classification label, the classification result is correct. For each kingdom dataset, we remove those groups with only one sequence and use both direct and weighted Euclidean metrics between k-mer NVs to get the corresponding 1NN accuracies (See Tables A4.1 to A4.7 of Appendix 4). The geometric metric with the highest 1NN accuracy is determined as the Natural Metric in the genome space. The Natural Metrics differ across various kingdoms, primarily due to variations in sequence distributions within each kingdom.

**Figure 3:**
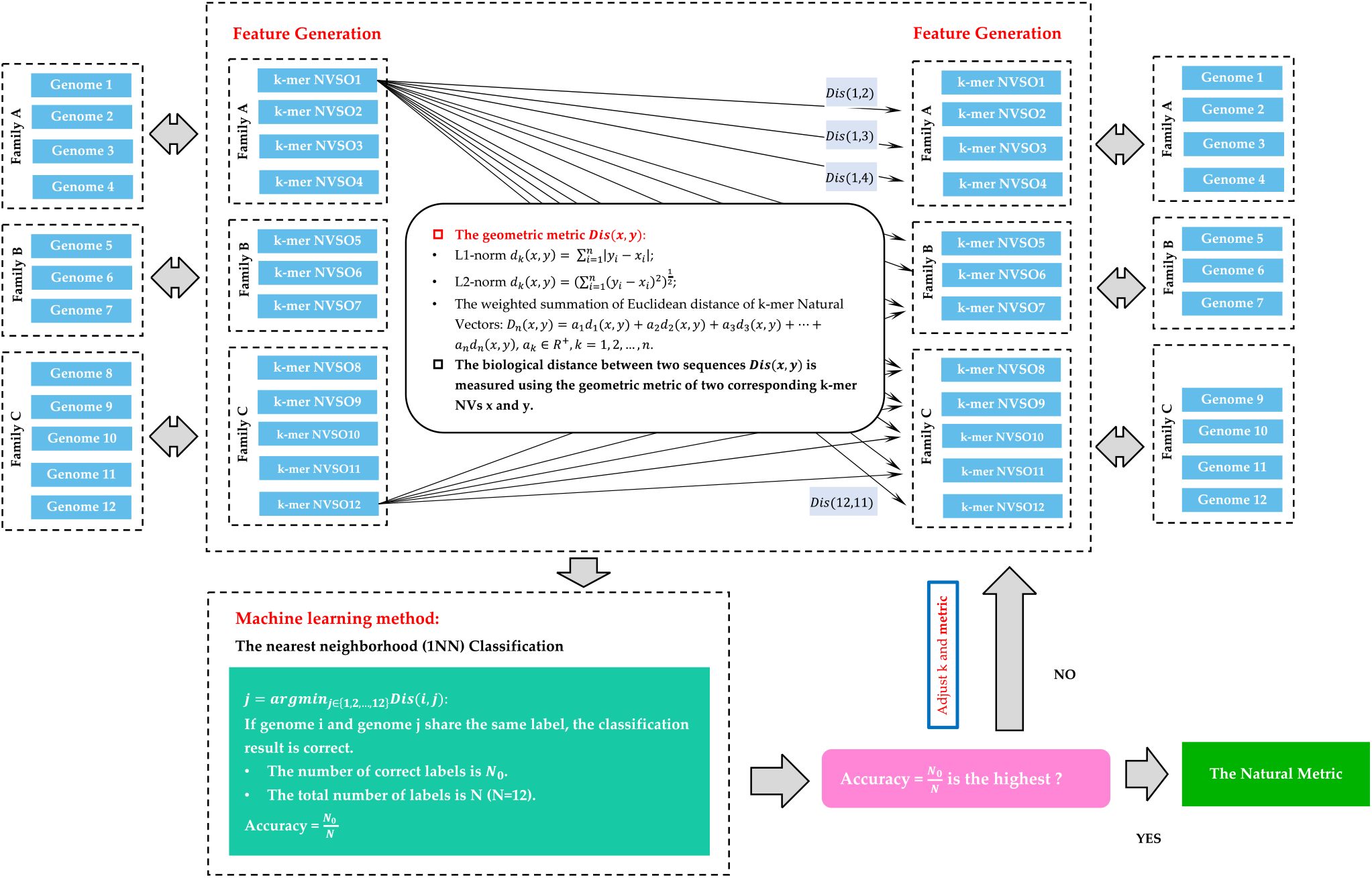
Flowchart of determining the Natural Metric. The metric is determined using 1NN method. The metric with the highest accuracy is proper in the genome space.

For instance, consider two pairs of sequences from different kingdoms. If the sequences in one kingdom are densely arranged compared to the other kingdom, it is possible that the pair of sequences within that kingdom appears closer in distance compared to the pair in the other kingdom. However, this does not necessarily imply a closer genetic relationship. It is important to emphasize that the concept of dense sequence arrangement we refer to here includes both detected and undetected sequences, analogous to dark matter.

We summarize the 1NN classification results in Column C of Table 1. For bacteria, fungi and plant datasets, the Natural Metric is determined as d_k_ (k = 9, 9, 2 respectively, L1-distance), and the corresponding maximum accuracies are 0.9030, 0.9056, and 0.8216, respectively. The nature metrics of the archaea, protozoa, vertebrate and invertebrate universe are 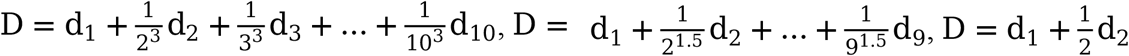, and 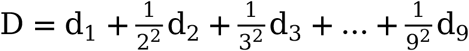, respectively (L1-distance). These metrics can well reflect the geometric relationship of the sequences, and can be used in phylogenetic analysis.

### 2.3. Geometric construction of Grand Biological Universe

Grand Biological Universe should contain all reliable biological sequences, and we try to find its geometric structure. We expect to reveal that the dimension of the Grand Biological Universe is the maximum of all various universe dimensions. In this Grand Universe, the universes of seven kingdoms are mutually disjoint, and the convex hull principle of genomes for each kingdom holds. In addition, the metric of the Grand Biological Universe should also be clarified, which can reflect the distance between two universes.

Now we determine the dimension of the Grand Biological Universe. From Column B of Table 1, the geometric structures of various universes exhibit existential discrepancies, that is, the NV dimensions are different when the convex hull principle holds. In order to accommodate all sequences of the seven kingdoms into the Grand Biological Universe and ensure that each small kingdom universe does not intersect each other in this grand universe, and that the convex hull principle of genomes holds in each small kingdom universe, we intuitively consider the maximum of all various universe dimensions. The NV corresponding to the spatial structure of the Grand Biological Universe is the combination of NV with 11-order moment and 2-mer NV with 2-order moment:

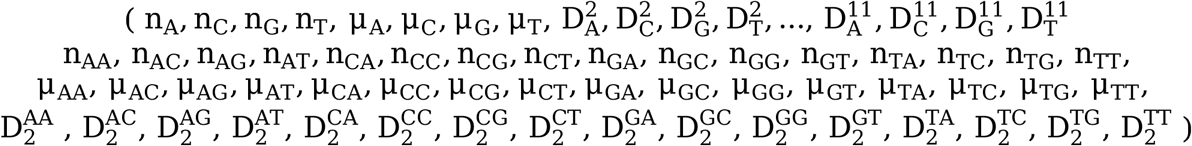

The vector is in the 96-dimensional Euclidean space, that is, the Grand Biological Universe is sitting in the 96-dimensional Euclidean space. Each sequence has a clear arrangement in this universe, and the 96-dimensional vector provides an advanced statistical description framework for nucleotide distribution.

The metric of the Grand Biological Universe should also be clear. This geometric metric can reflect the distance between two universes. We still use the 1NN method to test the metric, the correct classification criterion is that the two nearest sequences share the same kingdom label. The nearest neighborhood classification results (Figure 4) are based on 30121 sequences of seven kingdoms. The Grand Biological Universe is sitting in a 96-dimensional Euclidean space, and the corresponding 1NN classification accuracies are 0.9689 and 0.9666 for L1-distance and L2-distance, respectively. The accuracy is the greatest (0.9729) when the direct L1-distance d_2_ of 2-mer NVSOs, which indicates that the natural geometric metric is d_2_ in the Grand Biological Universe, and d_2_ can be used to measure the biological distance of each two universes.

**Figure 4:**
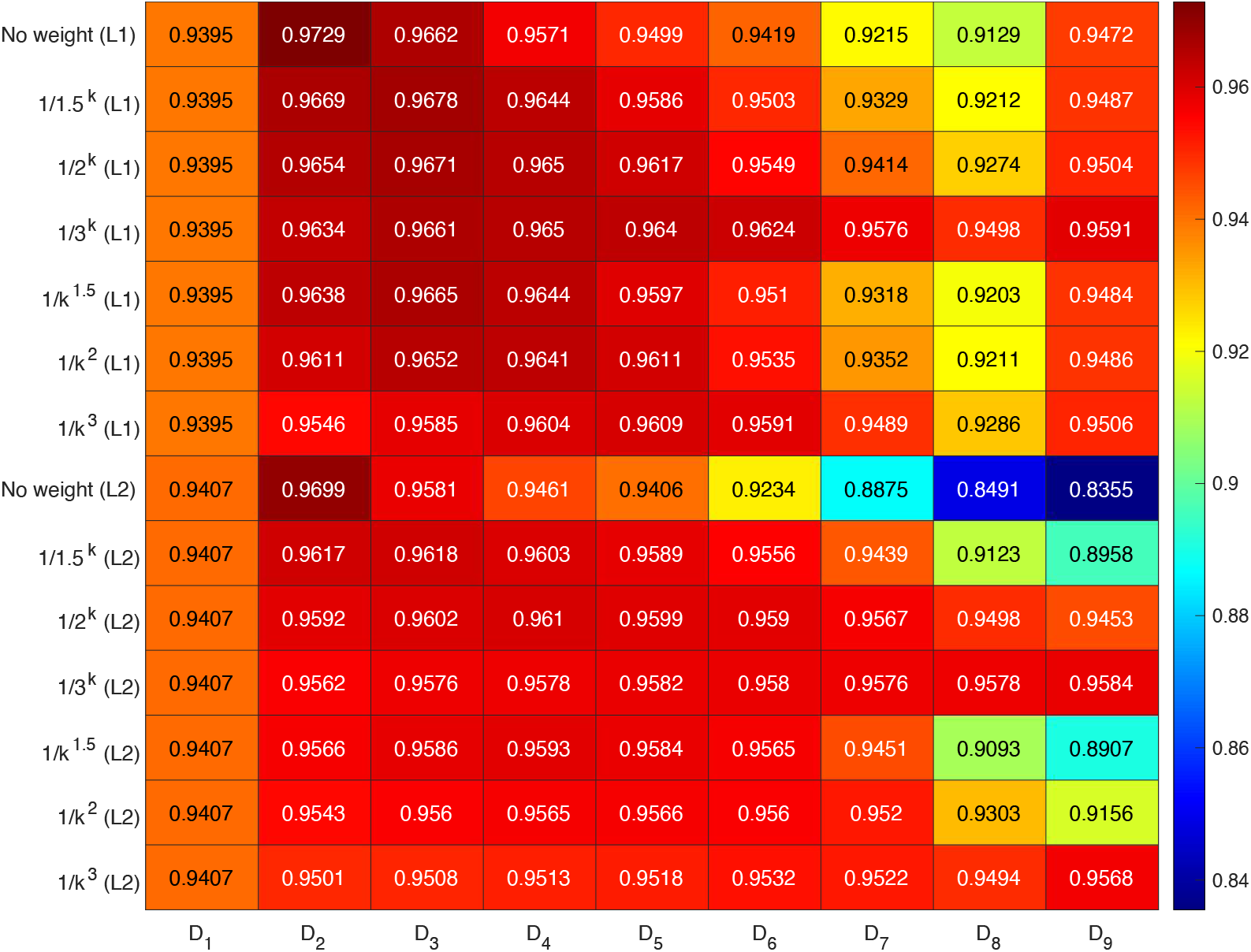
The Natural Metric determination in Grand Biological Universe. Accuracies under all metrics are over 0.8355, and the greatest value is 0.9729 when the metric is the direct L1-distance d_2_ of 2-mer NVSOs.

It should be noted that, according to our calculations, different universes in the Grand Biological Universe have different dimensions and measures. This suggests that attempts by traditional evolutionists to find an appropriate distance to measure the relationship between different species are likely to be futile. This is because different universes are filled with different dark matter, which distorts the time and space of biological universes to varying degrees.

### 2.4. An Application Example - Phylogenetic Analysis at the Family Level

Our natural vector approach has been demonstrated to be effective for phylogenetic analysis at the species level [4][15]. Here, in order to show the superiority and convenience of the grand biological universe, we conduct a phylogenetic analysis at the family level from the perspective of biological sequence. This is helpful to understand the relationship of high-order taxa from a sequence-based standpoint, especially for taxonomic groups with complex and controversial phylogeny, such as protozoa [25].

Specifically, in terms of methodology, we conduct phylogenetic analyses for bacteria and archaea using chromosomal sequences (genomes). For plants, protozoa, vertebrates, and invertebrates, we employ organelle sequences (chloroplasts for plants and mitochondria for other organisms) for phylogenetic inference. The detailed results can be found in Appendix 6, where we provide both the family names and their corresponding higher-order taxonomic units (note that for archaea, we only show the relationships at the phylum level) according to NCBI, aiding in the assessment of accuracy. We also show the results of Phaeophyceae protists and Reptiles as examples in Figure 5. Overall, our approach successfully revealed the relationships between different families, with most families belonging to the same higher-order taxonomic unit clustering together. Therefore, our phylogenetic analysis of high-order taxa obtained from the biological sequence perspective can provide new perspectives for taxonomists.

**Figure 5:**
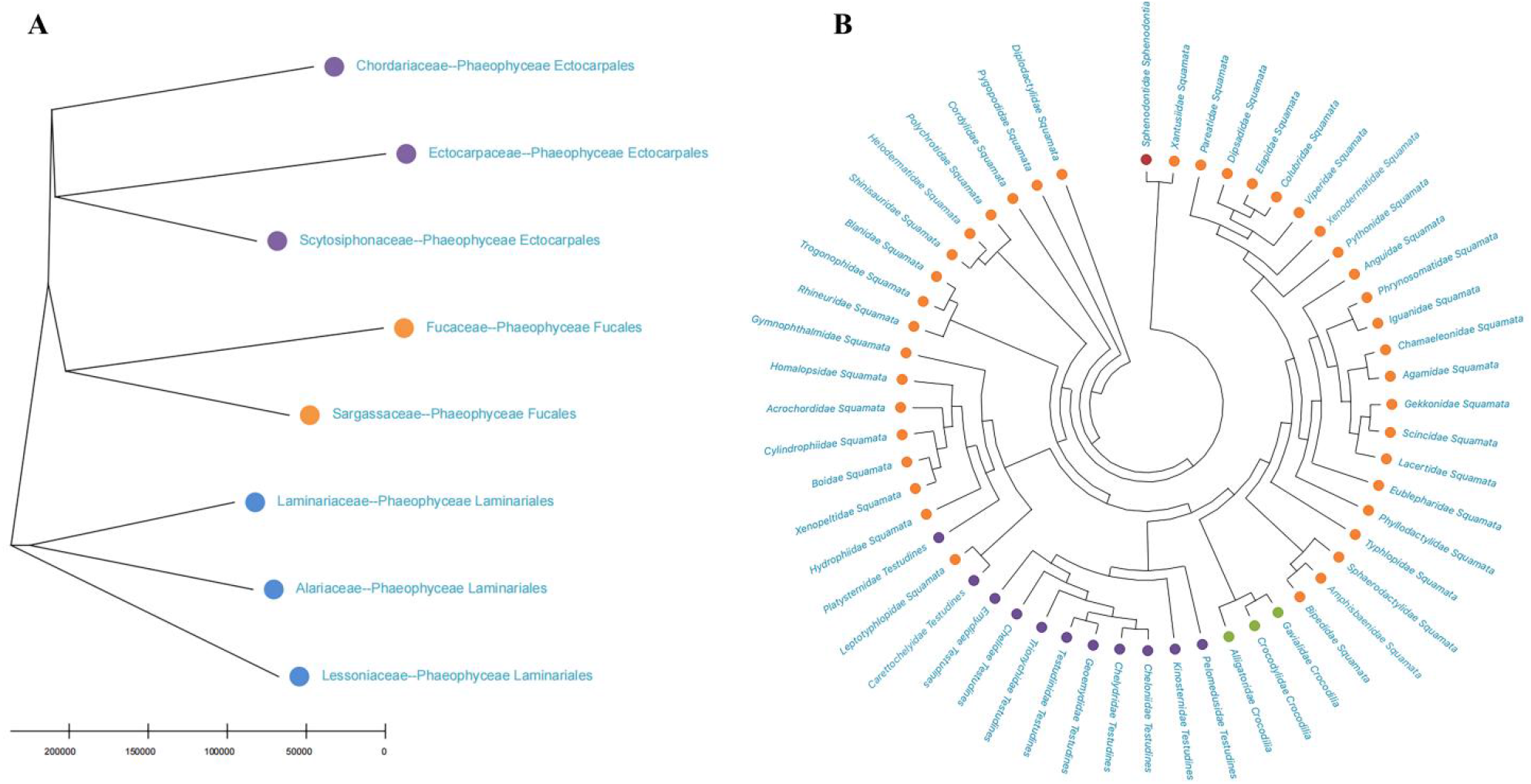
The phylogenetic results of Phaeophyceae protists (A) and Reptiles (B). The same color means the same higher-order taxonomic unit.

Performing phylogenetic analysis at the family level has revealed that most families belonging to the same higher-order taxonomic unit can be clustered together, thus validating the effectiveness of our constructed framework for the grand biological universe and the corresponding natural metric. This represents a novel perspective for classification research, where the relationships between high-order taxonomic units are explored based on the holistic nature of biological sequence spaces.

## 3. Discussion

The mathematical concept is used to study important problems in biology. The biological chromosome sequence is encoded as a k-mer Natural Vector and can be represented as a point in the Euclidean space, which indicates that the genome space can be canonically embedded into the high-dimensional Euclidean space. The k-mer NV incorporates statistical properties and can well characterize the biological feature of the genomic sequences. Sequences belonging to the same biological group distribute similarly, and the convex hull corresponding to each group can be constructed based on NVs of sequences. Then the convex hull principle of genomes is checked to determine the dimension of Euclidean space. The value of k and the central moment order of k-mer NVHO are capable to be adjusted to make the convex hull principle of genomes hold. To reveal the geometry of the genome space, we use the machine learning method (1NN) to find the proper Natural Metric, which can reflect the structural and functional relationship of genomes. The Natural Metric is actually the geometric metric, which satisfies the properties of symmetry, positive definiteness, and triangle inequality [19].

As applications of virus geometric space, researchers have explored the existing but undiscovered HIV sequence [20] or the early transmission of SARS-CoV-2 [19]. The convex hull principle of molecular biology points out that convex hulls of NVs corresponding to different biological groups do not overlap with each other. An established convex hull may have a large blank part, so there may be sequences with biological significance that have not been sequenced. These sequences may be mutated sequences or already exist but have not yet been discovered. An important problem is to find the points satisfying the condition in the convex hull. A genome sequence can be represented as a point corresponding to k-mer NV in a finite-dimensional space, and each biological group corresponds to a convex hull. Motivated by this, useful heuristic algorithms, such as Random-permutation Algorithm with Penalty (RAP) and Random-permutation Algorithm with Penalty and COs-trained Search (RAPCOS), have been developed [20]. In the past three years, COVID-19 used to spread around the world. Using the k-mer NN and Natural Metric, all reliable genomes of SARS-CoV-2 downloaded from GISAID are compared with RaTG13 and RmYN02, which are the bat-derived coronaviruses with genomes similar to SARS-CoV-2 [26][27]. The paper concludes that it is highly unlikely that China was the first country where the first human-to-human transmission of SARS-CoV-2 occurred [19]. These applications show the progressiveness of our theory.

The validation datasets have covered the chromosome sequences of all kingdoms of lives, which include bacteria, archaea, fungi, plants, protozoa, vertebrates, and invertebrates. We also test the theory based on organelle sequences for plants, protozoa, vertebrates, and invertebrates, as illustrated in Appendix 5. We take plants as an example to explain the results. There are 7150 non-redundant complete sequences of organelle. We deleted those sequences without family labels, and retained 7130 sequences belonging to 468 families. After discarding the sequences containing degenerate bases, there are 6240 sequences left, they belong to 439 families, of which there is only one sequence in 148 families. These genomes are from chloroplasts, plastids, and mitochondria, respectively. All convex hulls of families are disjoint in 24-dimensional space, and the metric is 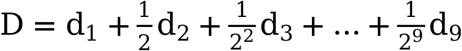. Our study is very meaningful since we have built a large database of k-mer NVs of seven kingdoms. The database is helpful for phylogeny and classification of genomic sequences, for example, microorganisms. Microorganisms are widely distributed, and mainly include bacteria, archaea, viruses, fungi, and some other protozoa. Archaea can live in extreme environments [28], and its classification still remains controversial. Protozoa are considered to be the simplest eukaryotes, which may cause diseases, such as malaria (by Plasmodium), Trichomoniasis (by Trichomonas vaginalis) and Amoebiasis (by Entamoeba histolytica) [29][30]. They occupy a key position in evolution and classification. We construct the genome universe for each kingdom, and the dimensions of embedded space exist discrepancy, which may hint at the relationship between genome space dimension and degree of evolution. As the degree of evolution increases, more representational dimensions may be required to truly capture the biological relationships of DNA sequences. In fact, our work has limitations. Currently, well-assembled sequencing data in NCBI is not enough, and the sequencing quality needs to be improved, which will affect our work. The more abundant the sequencing data and the higher the sequencing quality, the closer the genome space is to its real situation in nature. Therefore, we can construct a more complete genome space through this method in the future, which can better reflect the phylogeny and classification of biological sequences.

In addition, it should be noted that we conduct an application example of phylogenetic analysis at the family level. While overall our method effectively inferred relationships between different families, our results do not entirely align with the Taxonomy database [31] in NCBI. This discrepancy can be attributed, on one hand, to the incomplete sequencing data which may have influenced the range defined by convex hulls and consequently affected the clustering results. On the other hand, the assignment of many higher-level taxonomic units still remains controversial, such as Protozoa [25], Invertebrata [32], and Fungi [33], and our method purely relies on the analysis of biological sequences. In the future, as sequencing data continues to expand, the convex hulls will be filled and extended with newly acquired sequence data, thereby significantly improving the accuracy and rationality of genome space and natural metrics. This advancement will offer new perspectives for taxonomists. In addition, the grand biological universe and the corresponding natural metrics also help to outline the spatial structure of the genome/organelle sequences of different biological groups from a holistic perspective.

In conclusion, in previous study, we have demonstrated the convex hull principle of genomes and found the geometry of genome space for viruses [2], this paper further verified the theory for seven kingdoms. The datasets are well-assembled sequencing chromosome data downloaded from NCBI. Their corresponding genome space and Natural Metric are determined as follows: Bacterial genome space is sitting in 48-dimensional Euclidean space (NV with 11-order moment), and its metric is d_9_ (L1-distance). Archaea genome space is sitting in 48-dimensional Euclidean space (2-mer NV with 2-order moment), and its metric is D6_10_ (L1-distance). Fungi genome space is sitting in 40-dimensional Euclidean space (NV with 9-order moment), and its metric is d_9_ (L1-distance). Plant genome space is sitting in 24-dimensional Euclidean space (NV with 5-order moment), and its metric is d_2_ (L1-distance). Protozoa genome space is sitting in 36-dimensional Euclidean space (NV with 8-order moment), and its metric is D4_9_ (L1-distance). Vertebrate genome space is sitting in 28-dimensional Euclidean space (NV with 6-order moment), and its metric is D2_2_ (L1-distance). Invertebrate genome space is sitting in 28-dimensional Euclidean space (NV with 6-order moment), and its metric is D5_9_ (L1-distance). We also reveal the dimension and the corresponding metric of the Grand Biological Universe. The seven universes are mutually disjoint in 96-dimensional space, which also reveals that the dimension of the Grand Biological Universe is the maximum of all various universe dimensions. In addition, our findings demonstrate that diverse biological universes possess distinct dimensions and metrics, akin to the principles outlined in Einstein’s theory. In physics, space distortion is primarily attributed to the presence of dark matter. In a similar vein, we hypothesize that the distortion observed in the biological universe mainly arises from unobserved sequences. This observation inspires future research endeavors aimed at identifying unknown genomes through the exploration of such distortions. Finally, our work suggests that previous efforts to find a single distance to measure the relationship between all species are likely to be ineffective.

## 4. Materials and Methods

### 4.1. Method overview of sequence numerical representation

#### k-mer

If a genome sequence is S = s_1_s_2_s_3_…s_N_, s_i_ ∈ {A, C, G, T/U}, then a k-mer l_i_ is a possible substring of length k and the sequence S of length N has N-k+1 k-mers: L_1_ = s_1_…s_k_, L_2_ = s_2_…s_k+1_, …(L_N−k+1_ = s_N−k+1_…s_N_). For each given k, the number of k-mer is fixed. For example, 1-mers l_i_ (i = 1, 2, 3, 4) indicate A, C, G, T; 2-mers l_i_ (i = 1, 2, …, 16) include AA, AC, AG, AT, CA, CC, CG, CT, GA, GC, GG, GT, TA, TC, TG, TT; …; according to combinatorial mathematics, k-mers l_i_ (i = 1, 2, …, 4^k^) consist of 4^k^ subsequences in the case of DNA.

#### k-mer Natural Vector

Suppose that l_i_[j] is the indicator function of the j-th occurrence of a k-mer l_i_ in 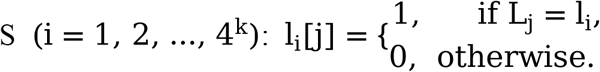 the distributions of a k-mer l_i_ can be described by three components:

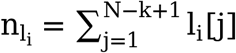denotes the counts of k-mer l_i_ in S;

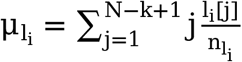 specifies the average location of k-mer l ;

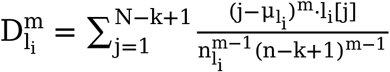 is the m-order central moment of emergence position of letter k-mer l_i (_m = 2, …, n, … .)

Then the k-mer Natural Vector with high order central moment (k-mer NVHO) for sequence S is defined by: 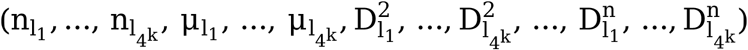. The dimension of central moment is 4^k^ • (n − 1), and the dimension of counts and average location of k-mers are both 4^k^, so the complete k-mer dimensional natural vector is 4^k^ • (n + 1) -dimensional (n = 2, 3, 4, …).

The k-mer Natural Vector with the second order central moment (k-mer NVSO) has been verified to be enough to represent the sequence and satisfy one-to-one mapping [4]. It is a 4^k^ · 3-dimensional vector:

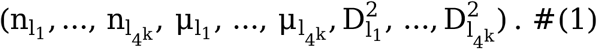

#### Natural Vector

If k in the k-mer Natural Vector is 1, the vector describes the distribution of the four nucleotides (A, C, G, T or U), and it is the traditional Natural Vector with high order central moment (NVHO), which is (4+4n)-dimensional (n=2, 3, 4, …):

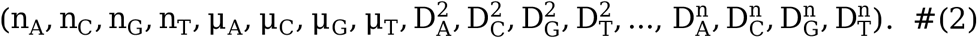

If we only consider the second central moment, the vector is traditional 12-dimensional natural vector with second central moment (NVSO): 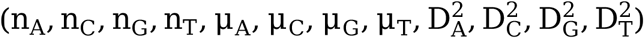.

An example. If the genomic sequence is CCGTAATCGTAG, the calculation example of the corresponding components of the k-mer NV can be found in Appendix 1.

### 4.2. The genome space construction

#### Convex hull concept in computational geometry

The convex hull of a finite point set C =x_1_, x_2_, …, x_k_, x_i_ ∈ R^n^ is the minimal convex set that contains these points [34]. Mathematically, it is the convex combinations of all points in C:

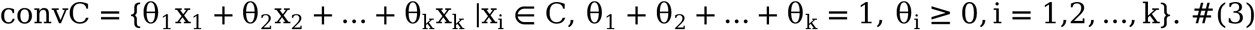

The convex hull is a convex polygon if all x_i_’s are 2-dimensional vectors, and convex polytope if all x_i_’s are high dimensional vectors.

#### Convex hull principle of genomes

In this study, x_i_ is the k-mer Natural Vector (k=1, 2, 3, …). Each sequence can be represented as a vector, and vectors from the same family form a convex hull. The convex hull principle of genomes states that convex hulls corresponding to different families are disjoint mutually [2].

Optimization method is used to check whether two convex hulls intersect. If A = Conv a_1_, a_2_, …, a_m_ and B = Conv b_1_, b_2_, …, b_n_ intersect, the convex combination of these points satisfy the formula: 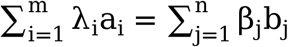, where 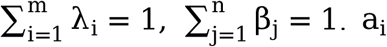 or b_j_ is the k-mer NV. It means that the following optimization problem has a feasible solution (That is, there are non-zero coefficients {λ_1_, λ_2_, …, λ_m_; β_1_, β_2_, …, β_n_} such that the minimum value of the following optimization problem is 0, otherwise the minimum value of the following optimization problem does not exist):

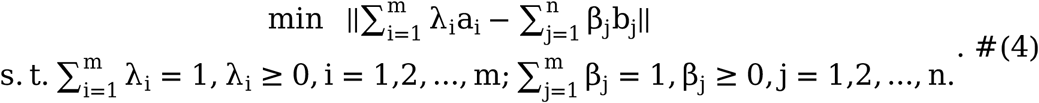

#### The genome space definition

The genome space contains all known genomes and reflects the important nature of the genome universe [2]. Mathematically, the genome space can be regarded as a moduli space and constructed as a subspace in a high-dimensional Euclidean space. If the convex hull principle holds in R^K^, the genome space exists and the genome space is located in a K-dimensional Euclidean space. Here K is the minimum dimension of the Euclidean space where the convex hull principle holds. The flowchart of constructing the genome space is illustrated in Figure 2.

### 4.3. Determination of Natural Metrics in genome space

#### The geometric metrics

The biological distance between two sequences Dis(x, y) is measured using the mathematical distance of their corresponding k-mer Natural Vectors: x and y, x, y ∈ R^n^. For example,

- L1-norm 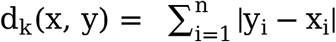;
- L2-norm 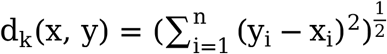;
- The weighted summation of Euclidean distance of k-mer Natural Vectors: D_n_(x, y) = a_1_d_1_(x, y) + a_2_d_2_(x, y) + a_3_d_3_(x, y) + … + a_n_d_n_ x, y, a_k_ ∈ R^+^, k = 1, 2, …, n. Here d_k_ is the L1-distance or L2-distance of k-mer Natural Vectors. The weight a_k_ reflects the contribution of the corresponding k-mer NV to the description of sequence similarity.

#### The nearest neighborhood (1NN) classification

The k-nearest neighbors classification algorithm (kNN) is a supervised machine learning method [35], and we only focus on k=1. The 1NN classification accuracy is used to determine the Natural Metric in genome space. For k-mer Natural Vector V1 of genome G1, we calculated another k-mer Natural Vector V2 of genome G2 nearest to V1. If the two corresponding genomes share the same label, the classification result is correct. The 1NN accuracy equals the number of correct labels N_0_ divided by the total number of labels 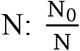. The geometric metric with the highest 1NN accuracy is the Natural Metric in the genome space. The flowchart of determining the Natural Metric is illustrated in Figure 3. The uncertainty of k gives the space to adjust the weights of D_n_ and improve the classification accuracy using the metric definition.

### 4.4. Biological sequence datasets analyzed

Understanding biological classification is necessary for our task, as shown in Figure 1. Based on Linnaean taxonomic ranks, the basic scheme of modern classification is developed, including domain, kingdom, phylum, class, order, family, genus, and species. The generally accepted domains include Archaea, Bacteria and Eukaryota, which is called three-domain system introduced by Carl Woese in 1977 [36][37]. In the meantime, all lives on earth can be divided into two parts according to whether they have the cellular structure: (1) Virus; (2) Two empires: Prokaryota and Eukaryota. Archaea and Bacteria can be merged into the Prokaryotes, and together with Fungi, Plants, Protozoa, Invertebrates and Vertebrates belonging to Eukaryotes, they form seven kingdoms [38]. Each kingdom has a more detailed classification rank: phylum, class, order, family, genus, etc. We use chromosome sequence datasets of the seven kingdoms to validate our genome universe theory.

The validation datasets are downloaded from the RefSeq database (https://ftp.ncbi.nlm.nih.gov/refseq/release/), Assembly GCF database (https://ftp.ncbi.nlm.nih.gov/genomes/refseq/), and Assembly GCA database (https://ftp.ncbi.nlm.nih.gov/genomes/genbank/) in March 2022. Reference sequences are usually used to answer the questions in the field of bioinformatics, GCF and GCA sequences were added to expand the dataset. The data are redundant, we removed three types of sequences: (1) unassembled sequences; (2) sequences with degenerate bases (Table A5.1 in Appendix 5); (3) sequences without classification labels, such as family. Only sequences with the header of “complete genome”-like or “chromosome”-like in the gbk file were retained. These data are reliable, and their summary information can be found in Column A of Table 1.

During data filtering, we found that each kind of data has its unique attributes. Let’s take the plant dataset as an example. In the three datasets (RefSeq; GCF; GCA), there are 8720 assembly chromosomal sequences, pertaining to 607 species and 90 families. Different species contain different numbers of chromosomes. Most of the chromosomal sequences are of low quality, and the number of chromosomal sequences without degenerate bases is 399, belonging to 22 families. The detailed description is provided in Appendix 2.

### 4.5. Phylogenetic Analysis of High-Order Taxonomic Units

In order to give an application example of the grand biological universe, we conduct a phylogenetic analysis at the family level from the perspective of biological sequence. During the phylogenetic analysis of high-order taxonomic units in each biological kingdom, we compute the average k-mer natural vectors of sequences in each family (in each phylum for Archaea). Subsequently, we utilize the Natural Metric of each kingdom to construct a distance matrix. Finally, FastME, a phylogenetic tree inference tool available at http://www.atgc-montpellier.fr/fastme/ [39], is used for tree construction.

## Acknowledgments

Professor Stephen S.-T. Yau is grateful to the National Center for Theoretical Sciences (NCTS) for providing an excellent research environment while part of this research was done.

## Author Contributions

Conceptualization: SSY

Methodology: SSY

Investigation: SSY, NS, HY, RR, TZ, MG, LZ

Visualization: NS, RR

Supervision: SSY

Writing—original draft: NS, RR

Writing—review & editing: SSY, NS, HY, RR, TZ, MG, LZ

## Data and materials availability

The data presented in this study can be downloaded in the public database, and also available in Supplementary Materials.

## Supplementary Materials

Appendix 1 to 6.

## Supplementary Materials for

### Other Supplementary Materials for this manuscript include the following

Data S1 to S3 (Appendix 2, 5, 6)

## Appendix 1 An example

Here we give an example on how to calculate the k-mer natural vector and natural vector (1)-(2) in the main text. If the genomic sequence is CCGTAATCGTAG, the corresponding components of vectors are calculated as follows:

### ▪ The example of 2-mer natural vector

**Table A1.1.** The j-th occurrence *l*_i_[*j*] of a 2-mer *l*_i_ (*l*_i_ = AA, AC, AG, AT, CA, CC, CG, CT, GA, GC, GG, GT, TA, TC, TG, TT) in CCGTAATCGTAG.

| Sliding window | CC | CG | GT | TA | AA | AT | TC | CG | GT | TA | AG | Count |
| --- | --- | --- | --- | --- | --- | --- | --- | --- | --- | --- | --- | --- |
|  | 1 | 2 | 3 | 4 | 5 | 6 | 7 | 8 | 9 | 10 | 11 |  |
| AA[j] |  |  |  |  | 1 |  |  |  |  |  |  | 1 |
| AC[j] |  |  |  |  |  |  |  |  |  |  |  | 0 |
| AG[j] |  |  |  |  |  |  |  |  |  |  | 1 | 1 |
| AT[j] |  |  |  |  |  | 1 |  |  |  |  |  | 1 |
| CA[j] |  |  |  |  |  |  |  |  |  |  |  | 0 |
| CC[j] | 1 |  |  |  |  |  |  |  |  |  |  | 1 |
| CG[j] |  | 1 |  |  |  |  |  | 1 |  |  |  | 2 |
| CT[j] |  |  |  |  |  |  |  |  |  |  |  | 0 |
| GA[j] |  |  |  |  |  |  |  |  |  |  |  | 0 |
| GC[j] |  |  |  |  |  |  |  |  |  |  |  | 0 |
| GG[j] |  |  |  |  |  |  |  |  |  |  |  | 0 |
| GT[j] |  |  | 1 |  |  |  |  |  | 1 |  |  | 2 |
| TA[j] |  |  |  | 1 |  |  |  |  |  | 1 |  | 2 |
| TC[j] |  |  |  |  |  |  | 1 |  |  |  |  | 1 |
| TG[j] |  |  |  |  |  |  |  |  |  |  |  | 0 |
| TT[j] |  |  |  |  |  |  |  |  |  |  |  | 0 |

1. 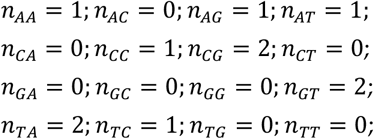
2. 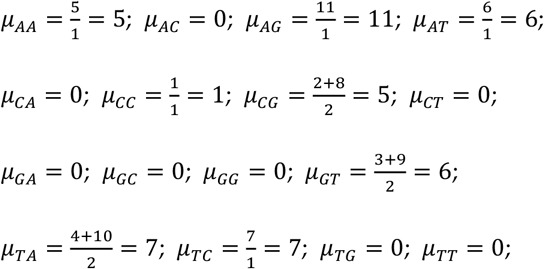
3. 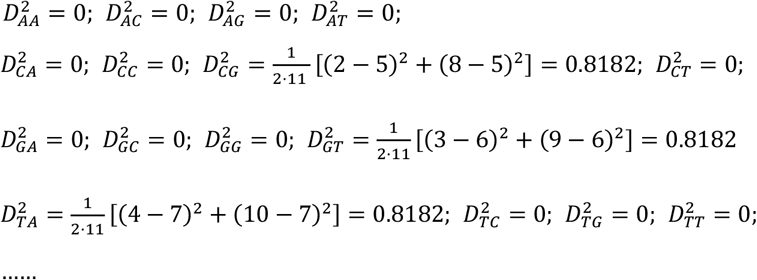
4. 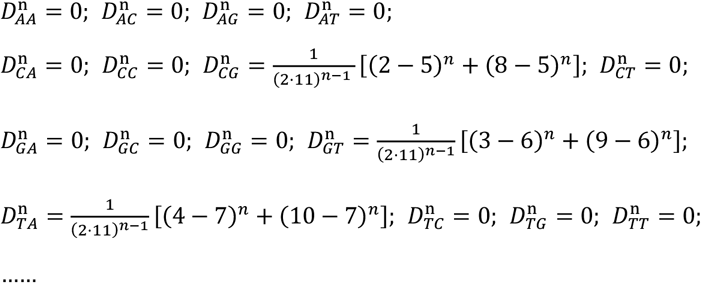

Thus, the 2-mer Natural Vector with high order central moment (2-mer NVHO) for sequence CCGTAATCGTAG is:

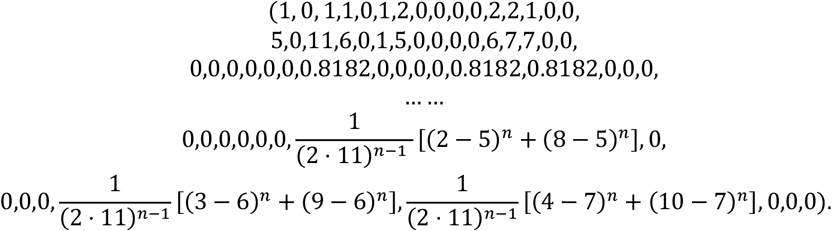

The 2-mer Natural Vector with second central moment (2-mer NVSO) corresponding to formula (1) is:

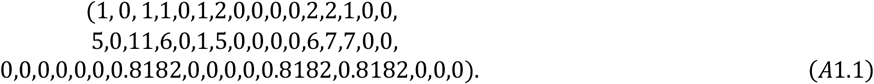

### ▪ The example of 1-mer natural vector

**Table A1.2.** The j-th occurrence *l*_i_[*j*] of a 1-mer *l*_i_ (*l*_i_ = A, C, G, T) in CCGTAATCGTAG.

|  | C | C | G | T | A | A | T | C | G | T | A | G | Count |
| --- | --- | --- | --- | --- | --- | --- | --- | --- | --- | --- | --- | --- | --- |
|  | 1 | 2 | 3 | 4 | 5 | 6 | 7 | 8 | 9 | 10 | 11 | 12 |  |
| A[j] |  |  |  |  | 1 | 1 |  |  |  |  | 1 |  | 3 |
| C[j] | 1 | 1 |  |  |  |  |  | 1 |  |  |  |  | 3 |
| G[j] |  |  | 1 |  |  |  |  |  | 1 |  |  | 1 | 3 |
| T[j] |  |  |  | 1 |  |  | 1 |  |  | 1 |  |  | 3 |

1. 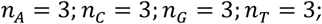
2. 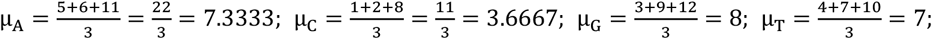
3. 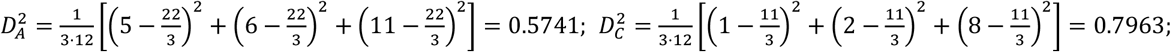

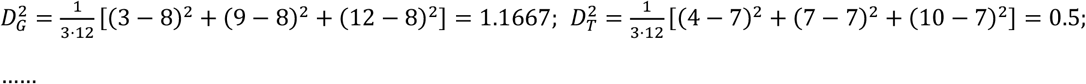
4. 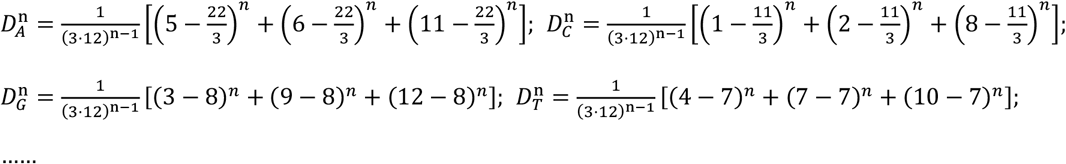

The Natural Vector with high order central moment (NVHO) is:

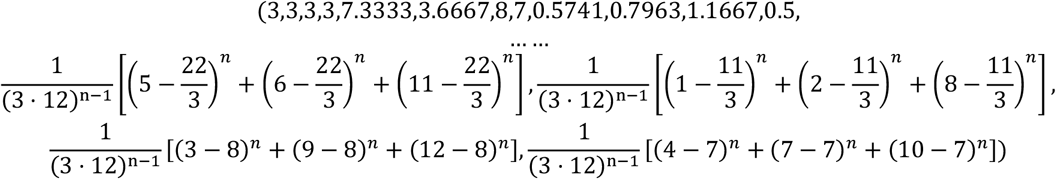

The traditional 12-dimensional Natural Vector with second order central moment (NVSO) corresponding to formula (2) is:

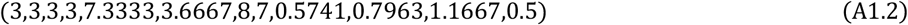

## Appendix 3: Convex Hull Analysis

Note:

### ▪ k-mer Natural Vector

The NVHO is 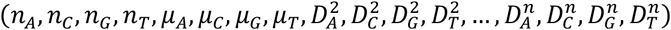, and it is in (4+4n)-dimensional Euclidean distance.

k-mer NVSO is 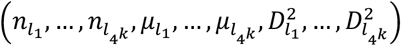, and it is in 4^*k*^ · 3-dimensional Euclidean distance.

### ▪ Convex hull concept in computational geometry

The convex hull of a finite point set *C* = {*x*_1_, *x*_2_, …, *x*_*k*_}, *x*_*i*_ ∈ *R*^*n*^ is the minimal convex set that contains these points. Mathematically, it is the convex combinations of all points in C:

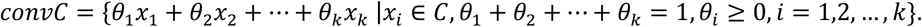

### ▪ Convex hull principle of genomes

In this study, *x*_*i*_ is the Natural Vector with high order central moment. Each sequence can be represented as a vector, and vectors from the same family form a convex hull. Convex hull principle of genomes states that convex hulls corresponding to different families are disjoint mutually.

### ▪ Optimization method is used to check whether two convex hulls intersect

If *A* = *Cov*{*a*_1_, *a*_2_, …, *a*_*m*_} and *B* = *Cov*{*b*_1_, *b*_2_, …, *b*_*n*_} intersect, the convex combination of these points satisfy the formula: 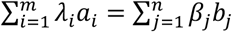 where 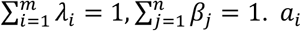. *a*_*i*_ or *b*_*j*_ is Natural Vector with high order central moment.

### ▪ The genome space definition

The genome space contains all known genomes and reflects the important nature of the genome universe. Mathematically, the genome space can be regarded as a moduli space and constructed as a subspace in a high-dimensional Euclidean space. If the convex hull principle holds in *R*^*K*^, the genome space exists and the genome space is located in a K-dimensional Euclidean space. Here K is the minimum dimension of the Euclidean space where the convex hull principle holds.

▪ The flowchart for constructing the genome space is illustrated in Figure 1.

**Table A3.1.**
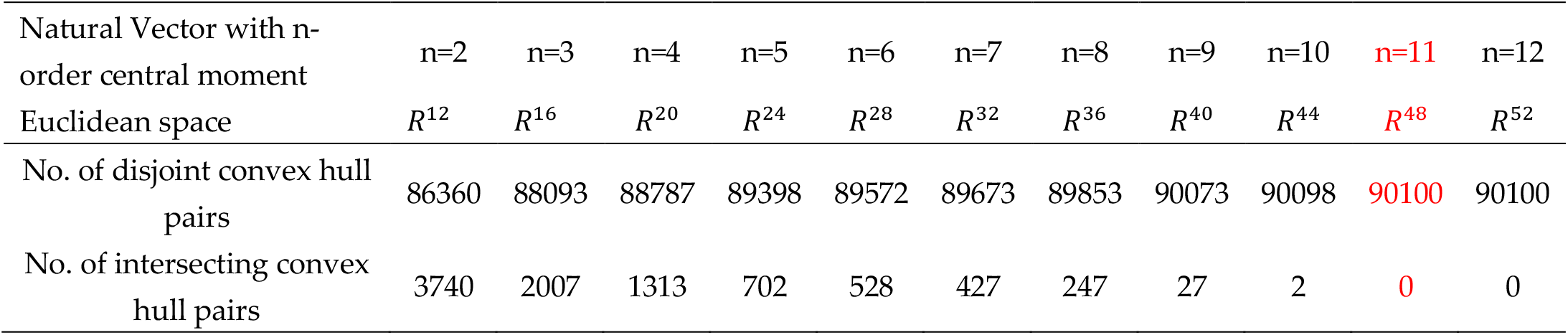
Convex hull analysis of Bacteria. The results are based on 24719 nucleoid sequences of 425 families. The number of disjoint convex hull pairs with the increase of the order of NVHO. There are 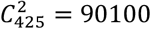 convex hull pairs totally, and all pairs are disjoint when the order is more than 11. According to the definition of embedding dimension of the moduli space, we choose the space with the lowest dimension, which indicates that the bacteria genome space is sitting in a 48-dimensional Euclidean space.

**Table A3.2.**
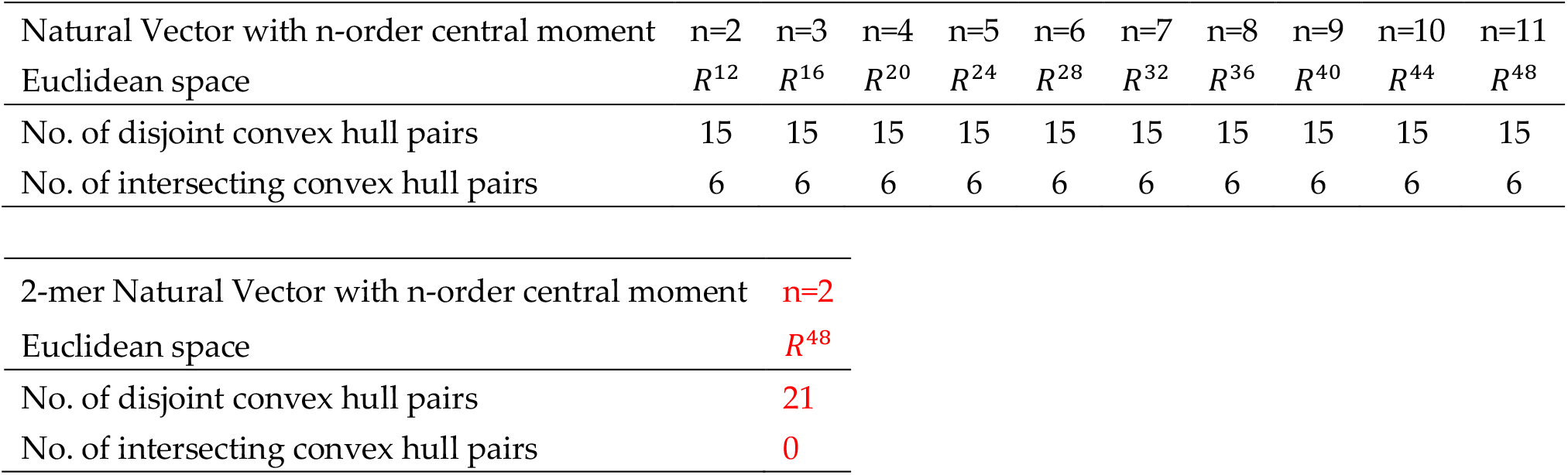
Convex hull analysis of Archaea. The results are based on 440 complete genomes of 7 phyla. There are 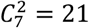 convex hull pairs totally, and all pairs are disjoint based on 2-mer Natural Vector with 2-order central moment. According to the definition of embedding dimension of the moduli space, we choose the space with the lowest dimension, which indicates that the archaea genome space is sitting in a 48-dimensional Euclidean space.

| Natural Vector with n-order central moment | n=2 | n=3 | n=4 | n=5 | n=6 | n=7 | n=8 | n=9 | n=10 | n=11 |
| --- | --- | --- | --- | --- | --- | --- | --- | --- | --- | --- |
| Euclidean space | $R^{12}$ | $R^{16}$ | $R^{20}$ | $R^{24}$ | $R^{28}$ | $R^{32}$ | $R^{36}$ | $R^{40}$ | $R^{44}$ | $R^{48}$ |
| No. of disjoint convex hull pairs | 15 | 15 | 15 | 15 | 15 | 15 | 15 | 15 | 15 | 15 |
| No. of intersecting convex hull pairs | 6 | 6 | 6 | 6 | 6 | 6 | 6 | 6 | 6 | 6 |

|  |  |
| --- | --- |
| 2-mer Natural Vector with n-order central moment | n=2 |
| Euclidean space | $R^{48}$ |
| No. of disjoint convex hull pairs | 21 |
| No. of intersecting convex hull pairs | 0 |

**Table A3.3.** Convex hull analysis of Fungi. The results are based on 2628 chromosome sequences of 23 families. The number of disjoint convex hull pairs with the increase of the order of NVHO. There are 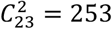 convex hull pairs totally, and all pairs are disjoint when the order is more than 9. According to the definition of embedding dimension of the moduli space, we choose the space with the lowest dimension, which indicates that the fungi genome space is sitting in a 40-dimensional Euclidean space.

| Natural Vector with n-order central moment | n=2 | n=3 | n=4 | n=5 | n=6 | n=7 | n=8 | n=9 | n=10 | n=11 |
| --- | --- | --- | --- | --- | --- | --- | --- | --- | --- | --- |
| Euclidean space | $R^{12}$ | $R^{16}$ | $R^{20}$ | $R^{24}$ | $R^{28}$ | $R^{32}$ | $R^{36}$ | $R^{40}$ | $R^{44}$ | $R^{48}$ |
| No. of disjoint convex hull pairs | 224 | 230 | 234 | 236 | 241 | 247 | 252 | 253 | 253 | 253 |
| No. of intersecting convex hull pairs | 29 | 23 | 19 | 17 | 12 | 6 | 1 | 0 | 0 | 0 |

**Table A3.4.** Convex hull analysis of Plant. The results are based on 399 chromosome sequences of 22 families. The number of disjoint convex hull pairs with the increase of the order of NVHO. There are 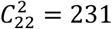 convex hull pairs totally, and all pairs are disjoint when the order is more than 5. According to the definition of embedding dimension of the moduli space, we choose the space with the lowest dimension, which indicates that the plant genome space is sitting in a 24-dimensional Euclidean space.

| Natural Vector with n-order central moment | n=2 | n=3 | n=4 | n=5 | n=6 | n=7 | n=8 | n=9 | n=10 | n=11 |
| --- | --- | --- | --- | --- | --- | --- | --- | --- | --- | --- |
| Euclidean space | $R^{12}$ | $R^{16}$ | $R^{20}$ | $R^{24}$ | $R^{28}$ | $R^{32}$ | $R^{36}$ | $R^{40}$ | $R^{44}$ | $R^{48}$ |
| No. of disjoint convex hull pairs | 230 | 230 | 230 | 231 | 231 | 231 | 231 | 231 | 231 | 231 |
| No. of intersecting convex hull pairs | 1 | 1 | 1 | 0 | 0 | 0 | 0 | 0 | 0 | 0 |

**Table A3.5.** Convex hull analysis of Protozoa. The results are based on 1200 chromosome sequences of 20 families. The number of disjoint convex hull pairs with the increase of the order of NVHO. There are 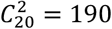 convex hull pairs totally, and all pairs are disjoint when the order is more than 8. According to the definition of embedding dimension of the moduli space, we choose the space with the lowest dimension, which indicates that the plant genome space is sitting in a 36-dimensional Euclidean space.

| Natural Vector with n-order central moment | n=2 | n=3 | n=4 | n=5 | n=6 | n=7 | n=8 | n=9 | n=10 |
| --- | --- | --- | --- | --- | --- | --- | --- | --- | --- |
| Euclidean space | $R^{12}$ | $R^{16}$ | $R^{20}$ | $R^{24}$ | $R^{28}$ | $R^{32}$ | $R^{36}$ | $R^{40}$ | $R^{44}$ |
| No. of disjoint convex hull pairs | 186 | 187 | 188 | 188 | 189 | 189 | 190 | 190 | 190 |
| No. of intersecting convex hull pairs | 4 | 3 | 2 | 2 | 1 | 1 | 0 | 0 | 0 |

**Table A3.6.** Convex hull analysis of Vertebrate. The results are based on 390 complete genomes of 4 classes. The number of disjoint convex hull pairs with the increase of the order of NVHO. There are 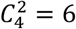 convex hull pairs totally, and all pairs are disjoint when the order is more than 6. According to the definition of embedding dimension of the moduli space, we choose the space with the lowest dimension, which indicates that the vertebrate genome space is sitting in a 28-dimensional Euclidean space.

| Natural Vector with n-order central moment | n=2 | n=3 | n=4 | n=5 | n=6 | n=7 | n=8 | n=9 | n=10 |
| --- | --- | --- | --- | --- | --- | --- | --- | --- | --- |
| Euclidean space | $R^{12}$ | $R^{16}$ | $R^{20}$ | $R^{24}$ | $R^{28}$ | $R^{32}$ | $R^{36}$ | $R^{40}$ | $R^{44}$ |
| No. of disjoint convex hull pairs | 3 | 3 | 4 | 4 | 6 | 6 | 6 | 6 | 6 |
| No. of intersecting convex hull pairs | 3 | 3 | 2 | 2 | 0 | 0 | 0 | 0 | 0 |

**Table A3.7.**
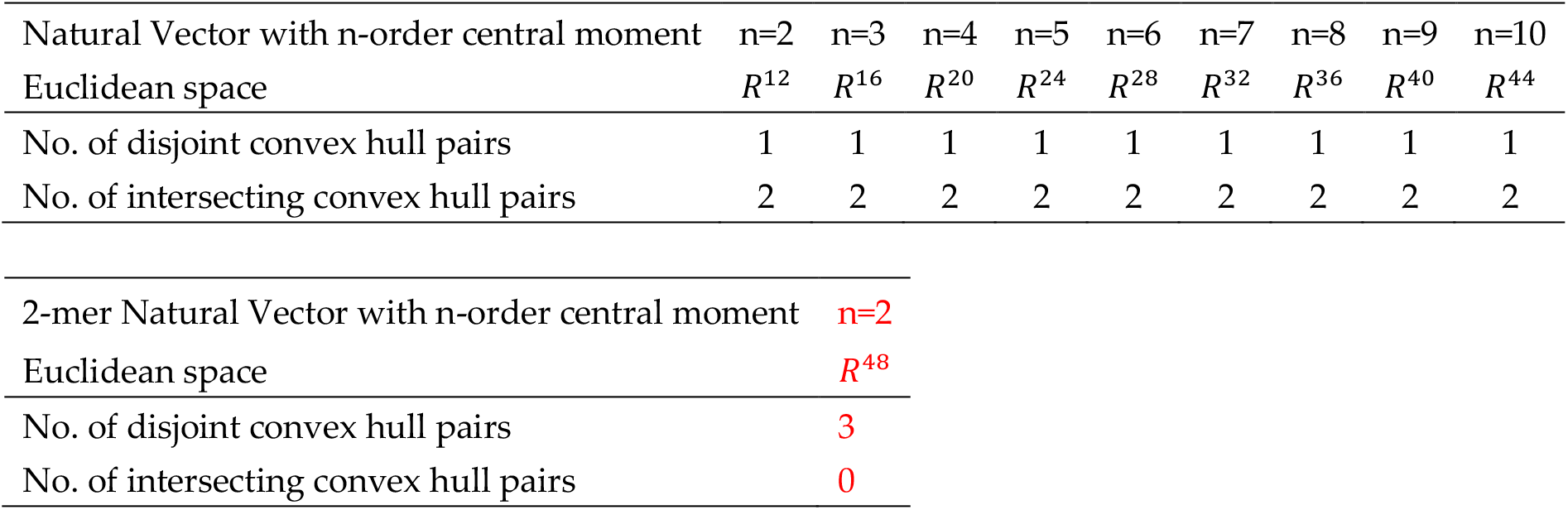
Convex hull analysis of Invertebrate. The results are based on 345 chromosome sequences of 3 orders. There are 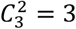 convex hull pairs totally, and all pairs are disjoint based on 2-mer Natural Vector with 2-order central moment. According to the definition of embedding dimension of the moduli space, we choose the space with the lowest dimension, which indicates that the invertebrate genome space is sitting in a 48-dimensional Euclidean space.

| Natural Vector with n-order central moment | n=2 | n=3 | n=4 | n=5 | n=6 | n=7 | n=8 | n=9 | n=10 |
| --- | --- | --- | --- | --- | --- | --- | --- | --- | --- |
| Euclidean space | $R^{12}$ | $R^{16}$ | $R^{20}$ | $R^{24}$ | $R^{28}$ | $R^{32}$ | $R^{36}$ | $R^{40}$ | $R^{44}$ |
| No. of disjoint convex hull pairs | 1 | 1 | 1 | 1 | 1 | 1 | 1 | 1 | 1 |
| No. of intersecting convex hull pairs | 2 | 2 | 2 | 2 | 2 | 2 | 2 | 2 | 2 |

|  |  |
| --- | --- |
| 2-mer Natural Vector with n-order central moment | **n=2** |
| Euclidean space | **$R^{48}$** |
| No. of disjoint convex hull pairs | **3** |
| No. of intersecting convex hull pairs | **0** |

## Appendix 4

### The Nearest Neighborhood Classification Results

Note:

### ▪ k-mer Natural Vector

The NVHO is 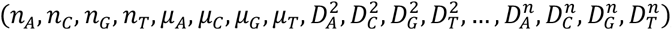, and it is in (4+4n)-dimensional Euclidean distance.

k-mer NVSO 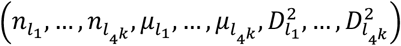, and it is in 4^*K*^ · 3-dimensional Euclidean distance.

### ▪ The geometric metrics

The biological distance between two sequences *d*_*k*_(*x, y*) is measured using the mathematical distance of their corresponding k-mer Natural Vectors: *x* and *y, x, y* ∈ *R*^*n*^. For example,

- L1-norm 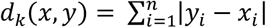;
- L2-norm 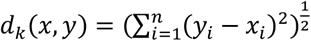;
- The weighted summation of Euclidean distance of k-mer Natural Vectors: *D*_*n*_(*x, y*) = *a*_1_*d*_1_(*x, y*) + *a*_2_*d*_2_(*x, y*) + *a*_3_*d*_3_(*x, y*) + … + *a*_*n*_*d*_*n*_(*x, y*), *a*_*k*_ ∈ *R, k* = 1, 2, …, *n*:
  a. If 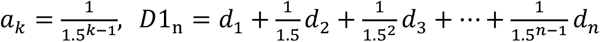
  b. If 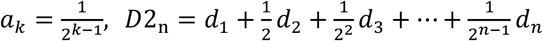
  c. If 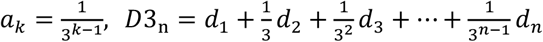
  d. If 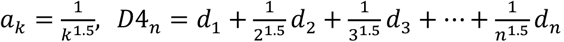
  e. If 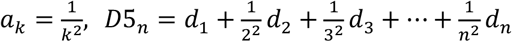
  f. If 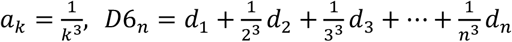

### ▪ The nearest neighborhood (1NN) classification

The 1NN classification accuracy is used to determine the geometric metric in genome space. For a genome G1, we calculated another genome G2 nearest to G1, if the two genomes have the same family label, the classification result is correct, and the 1NN accuracy equals the number of correct labels *N*_0_ divided by the total number of labels 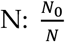. The metric with the highest 1NN accuracy is the natural metric in the genome space.

▪ The flowchart for determining the geometric metric is illustrated in **Figure 2** in the main text. The uncertainty of k gives the space to adjust the weights and improve the classification accuracy using the metric definition.

**Table A4.1.**
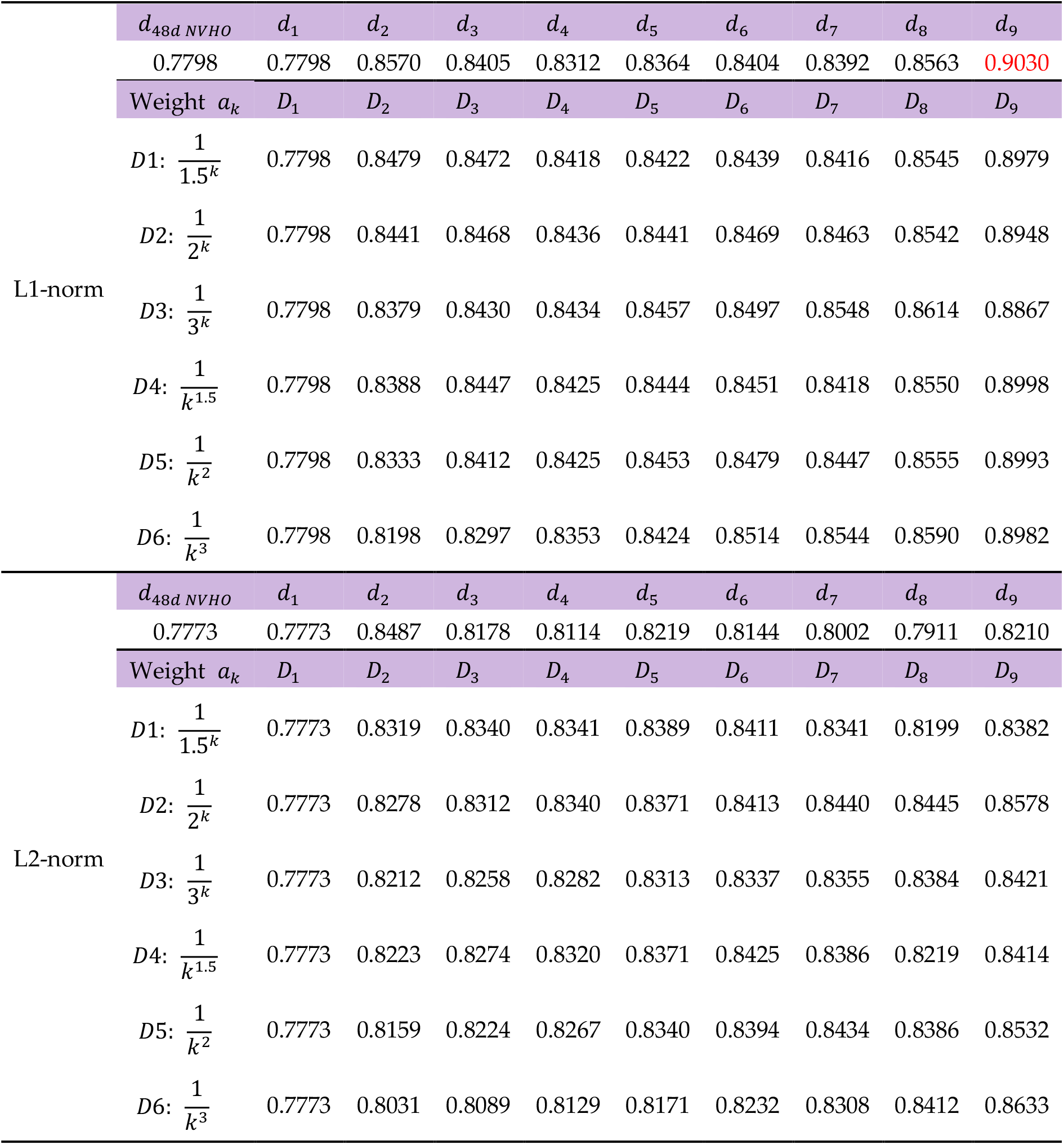
The nearest neighborhood classification accuracies of Bacteria family based on different metrics. The bacteria dataset includes 24719 nucleoid sequences belonging to 425 families, we removed those families with only one sequence, and the nearest neighborhood classification results are based on 24609 sequences of 315 families. The bacteria genome space is sitting in a 48-dimensional Euclidean space, and the corresponding 1NN classification accuracies are 0.7798 and 0.7773 for L1-distance and L2-distance, respectively. The accuracy is the greatest (0.9030) when the direct L1-distance *d*_9_ of 9-mer NVSOs, which indicates that the natural geometric metric is *d*_9_ in the bacteria genome space, and *d*_9_ can be used to measure the biological distance of two bacteria sequences.

**Table A4.2.** The nearest neighborhood classification accuracies of Archaea phyla based on different metrics. The archaea dataset includes 440 complete genomes belonging to 7 phyla, we removed those phyla with only one sequence, and the nearest neighborhood classification results are based on 438 complete genomes of 5 phyla. The archaea genome space is sitting in a 48-dimensional Euclidean space, and the corresponding 1NN classification accuracies are 0.8151 and 0.8036 for L1-distance and L2-distance, respectively. The accuracy is the greatest (0.9384) when the weighted L1-distance *D*6_10_ of 10-mer NVSOs, which indicates that the natural geometric metric is *D*6_10_ in the archaea genome space, and 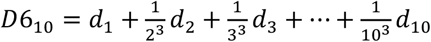 can be used to measure the biological distance of two archaea sequences.

| | $d_{48d\ 2merNVSO}$ | $d_1$ | $d_2$ | $d_3$ | $d_4$ | $d_5$ | $d_6$ | $d_7$ | $d_8$ | $d_9$ | $d_{10}$ |
| --- | --- | --- | --- | --- | --- | --- | --- | --- | --- | --- | --- |
|  | 0.8151 | 0.7352 | 0.8151 | 0.8151 | 0.8037 | 0.7945 | 0.7854 | 0.7534 | 0.7603 | 0.9064 | 0.9179 |
| | Weight $a_k$ | $D_1$ | $D_2$ | $D_3$ | $D_4$ | $D_5$ | $D_6$ | $D_7$ | $D_8$ | $D_9$ | $D_{10}$ |
| L1-norm | $D1: \frac{1}{1.5^k}$ | 0.7352 | 0.79 | 0.8037 | 0.7991 | 0.7968 | 0.7991 | 0.7763 | 0.7808 | 0.8858 | 0.9383 |
| | $D2: \frac{1}{2^k}$ | 0.7352 | 0.7877 | 0.7968 | 0.8059 | 0.7968 | 0.7922 | 0.7808 | 0.7854 | 0.8744 | 0.9338 |
| | $D3: \frac{1}{3^k}$ | 0.7352 | 0.7785 | 0.8014 | 0.7968 | 0.7991 | 0.7922 | 0.7854 | 0.7877 | 0.8653 | 0.9018 |
| | $D4: \frac{1}{k^{1.5}}$ | 0.7352 | 0.7785 | 0.7991 | 0.8105 | 0.79 | 0.7991 | 0.7694 | 0.7877 | 0.8927 | 0.9361 |
| | $D5: \frac{1}{k^2}$ | 0.7352 | 0.774 | 0.7991 | 0.8037 | 0.79 | 0.79 | 0.7763 | 0.7877 | 0.8927 | 0.9361 |
| | $D6: \frac{1}{k^3}$ | 0.7352 | 0.7557 | 0.7694 | 0.7808 | 0.79 | 0.7785 | 0.7717 | 0.7877 | 0.8836 | 0.9384 |
| | $d_{48d\ 2merNVSO}$ | $d_1$ | $d_2$ | $d_3$ | $d_4$ | $d_5$ | $d_6$ | $d_7$ | $d_8$ | $d_9$ | $d_{10}$ |
|  | 0.8036 | 0.7215 | 0.8036 | 0.8014 | 0.79 | 0.7831 | 0.7374 | 0.6621 | 0.5982 | 0.6689 | 0.8767 |
| | Weight $a_k$ | $D_1$ | $D_2$ | $D_3$ | $D_4$ | $D_5$ | $D_6$ | $D_7$ | $D_8$ | $D_9$ | $D_{10}$ |
| L2-norm | $D1: \frac{1}{1.5^k}$ | 0.7215 | 0.7854 | 0.7785 | 0.7945 | 0.7808 | 0.7717 | 0.774 | 0.7374 | 0.7397 | 0.8744 |
| | $D2: \frac{1}{2^k}$ | 0.7215 | 0.7763 | 0.774 | 0.7785 | 0.7831 | 0.7808 | 0.7831 | 0.7648 | 0.7808 | 0.8424 |
| | $D3: \frac{1}{3^k}$ | 0.7215 | 0.7694 | 0.774 | 0.7626 | 0.7626 | 0.7671 | 0.7626 | 0.7717 | 0.7763 | 0.7831 |
| | $D4: \frac{1}{k^{1.5}}$ | 0.7215 | 0.7694 | 0.7671 | 0.774 | 0.7854 | 0.7763 | 0.7694 | 0.7283 | 0.742 | 0.9041 |
| | $D5: \frac{1}{k^2}$ | 0.7215 | 0.7648 | 0.7717 | 0.7626 | 0.7763 | 0.774 | 0.7671 | 0.758 | 0.7717 | 0.8904 |
| | $D6: \frac{1}{k^3}$ | 0.7215 | 0.7466 | 0.7489 | 0.7557 | 0.7603 | 0.7626 | 0.758 | 0.7557 | 0.7763 | 0.8539 |

**Table A4.3.** The nearest neighborhood classification accuracies of Fungi family based on different metrics. The fungi dataset includes 2628 chromosome sequences belonging to 23 families, all families have more than two sequences, and the nearest neighborhood classification results are based on 2628 chromosome sequences of 23 families. The fungi genome space is sitting in a 40-dimensional Euclidean space, and the corresponding 1NN classification accuracies are 0.8607 and 0.8649 for L1-distance and L2-distance, respectively. The accuracy is the greatest (0.9056) when the direct L1-distance *d*_9_ of 9-mer NVSOs, which indicates that the natural geometric metric is *d*_9_ in the fungi genome space, and *d*_9_ can be used to measure the biological distance of two fungi sequences.

| | $d_{40d\ NVHO}$ | $d_1$ | $d_2$ | $d_3$ | $d_4$ | $d_5$ | $d_6$ | $d_7$ | $d_8$ | $d_9$ |
| --- | --- | --- | --- | --- | --- | --- | --- | --- | --- | --- |
|  | 0.8607 | 0.8607 | 0.8779 | 0.8569 | 0.8345 | 0.8307 | 0.8318 | 0.8322 | 0.8455 | 0.9056 |
| | Weight $a_k$ | $D_1$ | $D_2$ | $D_3$ | $D_4$ | $D_5$ | $D_6$ | $D_7$ | $D_8$ | $D_9$ |
| L1-norm | $D1: \frac{1}{1.5^k}$ | 0.8607 | 0.8771 | 0.8683 | 0.8596 | 0.8508 | 0.8375 | 0.8303 | 0.8436 | 0.8984 |
| | $D2: \frac{1}{2^k}$ | 0.8607 | 0.8756 | 0.8683 | 0.8630 | 0.8611 | 0.8535 | 0.8360 | 0.8379 | 0.8923 |
| | $D3: \frac{1}{3^k}$ | 0.8607 | 0.8744 | 0.8710 | 0.8661 | 0.8687 | 0.8702 | 0.8687 | 0.8569 | 0.8836 |
| | $D4: \frac{1}{k^{1.5}}$ | 0.8607 | 0.8756 | 0.8695 | 0.8623 | 0.8573 | 0.8402 | 0.8326 | 0.8451 | 0.9011 |
| | $D5: \frac{1}{k^2}$ | 0.8607 | 0.8710 | 0.8714 | 0.8653 | 0.8626 | 0.8539 | 0.8371 | 0.8451 | 0.9007 |
| | $D6: \frac{1}{k^3}$ | 0.8607 | 0.8691 | 0.8683 | 0.8657 | 0.8687 | 0.8744 | 0.8581 | 0.8425 | 0.8976 |
| | $d_{40d\ NVHO}$ | $d_1$ | $d_2$ | $d_3$ | $d_4$ | $d_5$ | $d_6$ | $d_7$ | $d_8$ | $d_9$ |
|  | 0.8649 | 0.8649 | 0.8714 | 0.8291 | 0.8033 | 0.8147 | 0.8151 | 0.8147 | 0.8250 | 0.8520 |
| | Weight $a_k$ | $D_1$ | $D_2$ | $D_3$ | $D_4$ | $D_5$ | $D_6$ | $D_7$ | $D_8$ | $D_9$ |
| L2-norm | $D1: \frac{1}{1.5^k}$ | 0.8649 | 0.8718 | 0.8607 | 0.8588 | 0.8581 | 0.8615 | 0.8463 | 0.8166 | 0.8402 |
| | $D2: \frac{1}{2^k}$ | 0.8649 | 0.8725 | 0.8653 | 0.8623 | 0.8626 | 0.8634 | 0.8672 | 0.8611 | 0.8642 |
| | $D3: \frac{1}{3^k}$ | 0.8649 | 0.8702 | 0.8687 | 0.8664 | 0.8672 | 0.8699 | 0.8702 | 0.8710 | 0.8721 |
| | $D4: \frac{1}{k^{1.5}}$ | 0.8649 | 0.8706 | 0.8661 | 0.8623 | 0.8611 | 0.8630 | 0.8573 | 0.8231 | 0.8463 |
| | $D5: \frac{1}{k^2}$ | 0.8649 | 0.8687 | 0.8687 | 0.8653 | 0.8649 | 0.8691 | 0.8702 | 0.8531 | 0.8558 |
| | $D6: \frac{1}{k^3}$ | 0.8649 | 0.8672 | 0.8657 | 0.8687 | 0.8699 | 0.8710 | 0.8721 | 0.8725 | 0.8782 |

**Table A4.4.** The nearest neighborhood classification accuracies of Plant family based on different metrics. The plant dataset includes 399 chromosome sequences belonging to 22 families, we removed those families with only one sequence, and the nearest neighborhood classification results are based on 398 chromosome sequences of 21 families. The plant genome space is sitting in a 24-dimensional Euclidean space, and the corresponding 1NN classification accuracies are 0.7915 and 0.7965 for L1-distance and L2-distance, respectively. The accuracy is the greatest (0.8216) when the direct L1-distance *d*_2_ of 2-mer NVSOs, which indicates that the natural geometric metric is *d*_2_ in the plant genome space, and *d*_2_ can be used to measure the biological distance of two plant sequences.

| | $d_{24d\ NVHO}$ | $d_1$ | $d_2$ | $d_3$ | $d_4$ | $d_5$ | $d_6$ | $d_7$ | $d_8$ | $d_9$ |
| --- | --- | --- | --- | --- | --- | --- | --- | --- | --- | --- |
|  | 0.7915 | 0.7915 | 0.8216 | 0.7663 | 0.7412 | 0.7538 | 0.7638 | 0.7839 | 0.7638 | 0.8216 |
| | Weight $a_k$ | $D_1$ | $D_2$ | $D_3$ | $D_4$ | $D_5$ | $D_6$ | $D_7$ | $D_8$ | $D_9$ |
| L1-norm | $D1: \frac{1}{1.5^k}$ | 0.7915 | 0.8191 | 0.8040 | 0.7764 | 0.7638 | 0.7688 | 0.7739 | 0.7789 | 0.8090 |
| | $D2: \frac{1}{2^k}$ | 0.7915 | 0.8166 | 0.8090 | 0.7789 | 0.7688 | 0.7688 | 0.7789 | 0.7839 | 0.8090 |
| | $D3: \frac{1}{3^k}$ | 0.7915 | 0.8090 | 0.8116 | 0.8040 | 0.7839 | 0.7864 | 0.7839 | 0.7915 | 0.8040 |
| | $D4: \frac{1}{k^{1.5}}$ | 0.7915 | 0.8116 | 0.8116 | 0.7764 | 0.7663 | 0.7714 | 0.7739 | 0.7688 | 0.8040 |
| | $D5: \frac{1}{k^2}$ | 0.7915 | 0.8065 | 0.8065 | 0.7940 | 0.7739 | 0.7688 | 0.7764 | 0.7764 | 0.8040 |
| | $D6: \frac{1}{k^3}$ | 0.7915 | 0.8015 | 0.8065 | 0.8015 | 0.7940 | 0.7889 | 0.7839 | 0.7864 | 0.8216 |
| | $d_{24d\ NVHO}$ | $d_1$ | $d_2$ | $d_3$ | $d_4$ | $d_5$ | $d_6$ | $d_7$ | $d_8$ | $d_9$ |
|  | 0.7965 | 0.7965 | 0.7965 | 0.7437 | 0.7211 | 0.7412 | 0.7588 | 0.7563 | 0.7462 | 0.7638 |
| | Weight $a_k$ | $D_1$ | $D_2$ | $D_3$ | $D_4$ | $D_5$ | $D_6$ | $D_7$ | $D_8$ | $D_9$ |
| L2-norm | $D1: \frac{1}{1.5^k}$ | 0.7965 | 0.8141 | 0.8040 | 0.7864 | 0.7714 | 0.7764 | 0.7789 | 0.7764 | 0.7638 |
| | $D2: \frac{1}{2^k}$ | 0.7965 | 0.8141 | 0.8090 | 0.7965 | 0.7839 | 0.7839 | 0.7889 | 0.7915 | 0.7990 |
| | $D3: \frac{1}{3^k}$ | 0.7965 | 0.8116 | 0.8040 | 0.7990 | 0.8015 | 0.8040 | 0.8040 | 0.8040 | 0.8040 |
| | $D4: \frac{1}{k^{1.5}}$ | 0.7965 | 0.8116 | 0.8040 | 0.7965 | 0.7839 | 0.7839 | 0.7814 | 0.7764 | 0.7864 |
| | $D5: \frac{1}{k^2}$ | 0.7965 | 0.8116 | 0.8040 | 0.7965 | 0.7965 | 0.7940 | 0.7990 | 0.7990 | 0.7965 |
| | $D6: \frac{1}{k^3}$ | 0.7965 | 0.8090 | 0.8065 | 0.8040 | 0.8015 | 0.7990 | 0.7965 | 0.8015 | 0.8090 |

**Table A4.5.**
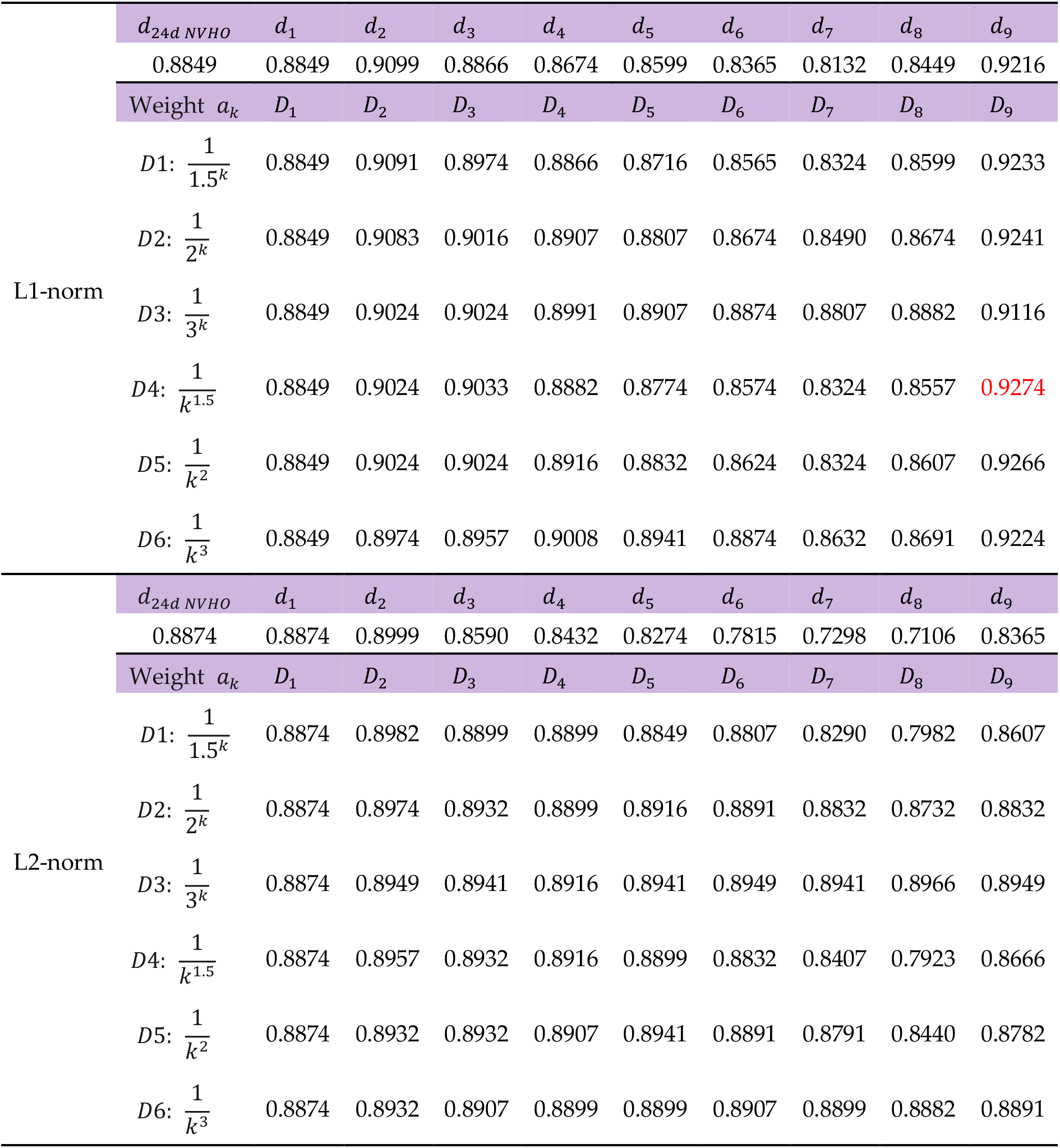
The nearest neighborhood classification accuracies of Protozoa family based on different metrics. The protozoa dataset includes 1200 chromosome sequences belonging to 20 families, we removed those families with only one sequence, and the nearest neighborhood classification results are based on 1199 chromosome sequences of 19 families. The protozoa genome space is sitting in a 36-dimensional Euclidean space, and the corresponding 1NN classification accuracies are both 0.8849 and 0.8874 for L1-distance and L2-distance, respectively. The accuracy is the greatest (0.9274) when the weighted L1-distance *D*4_9_ of 9-mer NVSOs, which indicates that the natural geometric metric is 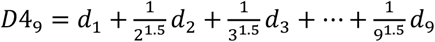 in the protozoa genome space, and *D*4_9_ can be used to measure the biological distance of two protozoa sequences.

**Table A4.6.** The nearest neighborhood classification accuracies of Vertebrate class based on different metrics. The vertebrate dataset includes 390 chromosome sequences belonging to 4 classes, all classes have more than two sequences, and the nearest neighborhood classification results are based on 390 chromosome sequences of 4 classes. The vertebrate genome space is sitting in a 28-dimensional Euclidean space, and the corresponding 1NN classification accuracies are both 0.8256 for L1-distance and L2-distance. The accuracy is the greatest (0.8590) when the weighted L1-distance *D*2_2_ of 2-mer NVSOs, which indicates that the natural geometric metric is 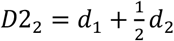 in the vertebrate genome space, and *D*2_2_ can be used to measure the biological distance of two vertebrate sequences.

| | $d_{28d\ NVHO}$ | $d_1$ | $d_2$ | $d_3$ | $d_4$ | $d_5$ | $d_6$ | $d_7$ | $d_8$ | $d_9$ |
| --- | --- | --- | --- | --- | --- | --- | --- | --- | --- | --- |
|  | 0.8256 | 0.8256 | 0.8513 | 0.8231 | 0.8077 | 0.8128 | 0.8359 | 0.8385 | 0.8308 | 0.8000 |
| | Weight $a_k$ | $D_1$ | $D_2$ | $D_3$ | $D_4$ | $D_5$ | $D_6$ | $D_7$ | $D_8$ | $D_9$ |
| L1-norm | $D1: \frac{1}{1.5^k}$ | 0.8256 | 0.8590 | 0.841 | 0.8205 | 0.8103 | 0.8231 | 0.8333 | 0.8282 | 0.8026 |
| | $D2: \frac{1}{2^k}$ | 0.8256 | 0.8590 | 0.8462 | 0.8231 | 0.8128 | 0.8128 | 0.8308 | 0.8333 | 0.8154 |
| | $D3: \frac{1}{3^k}$ | 0.8256 | 0.8487 | 0.8436 | 0.8333 | 0.8231 | 0.8179 | 0.8231 | 0.8359 | 0.8308 |
| | $D4: \frac{1}{k^{1.5}}$ | 0.8256 | 0.8538 | 0.8462 | 0.8256 | 0.8128 | 0.8231 | 0.8333 | 0.8231 | 0.7974 |
| | $D5: \frac{1}{k^2}$ | 0.8256 | 0.8436 | 0.8436 | 0.8282 | 0.8154 | 0.8179 | 0.8333 | 0.8282 | 0.7974 |
| | $D6: \frac{1}{k^3}$ | 0.8256 | 0.8359 | 0.841 | 0.841 | 0.8282 | 0.8231 | 0.8359 | 0.8359 | 0.8026 |
| | $d_{28d\ NVHO}$ | $d_1$ | $d_2$ | $d_3$ | $d_4$ | $d_5$ | $d_6$ | $d_7$ | $d_8$ | $d_9$ |
|  | 0.8256 | 0.8256 | 0.8282 | 0.7897 | 0.7974 | 0.8205 | 0.8282 | 0.8256 | 0.7923 | 0.6692 |
| | Weight $a_k$ | $D_1$ | $D_2$ | $D_3$ | $D_4$ | $D_5$ | $D_6$ | $D_7$ | $D_8$ | $D_9$ |
| L2-norm | $D1: \frac{1}{1.5^k}$ | 0.8256 | 0.8231 | 0.8077 | 0.8077 | 0.8128 | 0.8256 | 0.8256 | 0.8205 | 0.7872 |
| | $D2: \frac{1}{2^k}$ | 0.8256 | 0.8282 | 0.8205 | 0.8128 | 0.8179 | 0.8231 | 0.8256 | 0.8256 | 0.8179 |
| | $D3: \frac{1}{3^k}$ | 0.8256 | 0.8256 | 0.8231 | 0.8256 | 0.8205 | 0.8205 | 0.8179 | 0.8205 | 0.8154 |
| | $D4: \frac{1}{k^{1.5}}$ | 0.8256 | 0.8308 | 0.8282 | 0.8179 | 0.8179 | 0.8308 | 0.8205 | 0.8179 | 0.7769 |
| | $D5: \frac{1}{k^2}$ | 0.8256 | 0.8282 | 0.8231 | 0.8231 | 0.8205 | 0.8256 | 0.8333 | 0.8205 | 0.7923 |
| | $D6: \frac{1}{k^3}$ | 0.8256 | 0.8333 | 0.8256 | 0.8282 | 0.8231 | 0.8231 | 0.8333 | 0.8359 | 0.8205 |

**Table A4.7.** The nearest neighborhood classification accuracies of Invertebrates family based on different metrics. The invertebrate dataset includes 345 chromosome sequences belonging to 3 orders: Hymenoptera, Lepidoptera and Diptera, all orders have more than two sequences, and the nearest neighborhood classification results are based on 345 chromosome sequences of 3 orders. The invertebrate genome space is sitting in a 48-dimensional Euclidean space, and the corresponding 1NN classification accuracies are 0.8275 and 0.8609 for L1-distance and L2-distance, respectively. The accuracy is the greatest (0.8783) when the weighted L1-distance *D*5_9_ of 9-mer NVSOs, which indicates that the natural geometric metric is 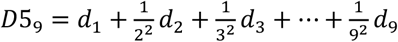 in the invertebrate genome space, and *D*5_9_ can be used to measure the biological distance of two invertebrate sequences.

| | $d_{48d\ 2merNVSO}$ | $d_1$ | $d_2$ | $d_3$ | $d_4$ | $d_5$ | $d_6$ | $d_7$ | $d_8$ | $d_9$ |
| --- | --- | --- | --- | --- | --- | --- | --- | --- | --- | --- |
|  | 0.8275 | 0.8319 | 0.8275 | 0.8522 | 0.8435 | 0.8493 | 0.8609 | 0.8667 | 0.8725 | 0.8696 |
| | Weight $a_k$ | $D_1$ | $D_2$ | $D_3$ | $D_4$ | $D_5$ | $D_6$ | $D_7$ | $D_8$ | $D_9$ |
| L1-norm | $D1: \frac{1}{1.5^k}$ | 0.8319 | 0.8696 | 0.8522 | 0.8493 | 0.8493 | 0.8638 | 0.8667 | 0.8667 | 0.8783 |
| | $D2: \frac{1}{2^k}$ | 0.8319 | 0.8638 | 0.8609 | 0.8493 | 0.8522 | 0.8609 | 0.8638 | 0.8638 | 0.8725 |
| | $D3: \frac{1}{3^k}$ | 0.8319 | 0.8638 | 0.8609 | 0.8493 | 0.8551 | 0.8551 | 0.8609 | 0.8609 | 0.8754 |
| | $D4: \frac{1}{k^{1.5}}$ | 0.8319 | 0.8638 | 0.8551 | 0.8493 | 0.8522 | 0.8638 | 0.8667 | 0.8667 | 0.8783 |
| | $D5: \frac{1}{k^2}$ | 0.8319 | 0.8609 | 0.8580 | 0.8493 | 0.8493 | 0.8609 | 0.8667 | 0.8667 | 0.8783 |
| | $D6: \frac{1}{k^3}$ | 0.8319 | 0.8522 | 0.8551 | 0.8580 | 0.8493 | 0.8609 | 0.8667 | 0.8667 | 0.8725 |
| | $d_{48d\ 2merNVSO}$ | $d_1$ | $d_2$ | $d_3$ | $d_4$ | $d_5$ | $d_6$ | $d_7$ | $d_8$ | $d_9$ |
|  | 0.8609 | 0.8261 | 0.8609 | 0.8348 | 0.8319 | 0.8377 | 0.8406 | 0.8435 | 0.8493 | 0.8435 |
| | Weight $a_k$ | $D_1$ | $D_2$ | $D_3$ | $D_4$ | $D_5$ | $D_6$ | $D_7$ | $D_8$ | $D_9$ |
| L2-norm | $D1: \frac{1}{1.5^k}$ | 0.8261 | 0.8580 | 0.8493 | 0.8406 | 0.8377 | 0.8377 | 0.8493 | 0.8609 | 0.8580 |
| | $D2: \frac{1}{2^k}$ | 0.8261 | 0.8580 | 0.8522 | 0.8493 | 0.8435 | 0.8435 | 0.8493 | 0.8580 | 0.8580 |
| | $D3: \frac{1}{3^k}$ | 0.8261 | 0.8609 | 0.8522 | 0.8464 | 0.8464 | 0.8435 | 0.8406 | 0.8435 | 0.8493 |
| | $D4: \frac{1}{k^{1.5}}$ | 0.8261 | 0.8609 | 0.8493 | 0.8377 | 0.8377 | 0.8406 | 0.8522 | 0.8580 | 0.8464 |
| | $D5: \frac{1}{k^2}$ | 0.8261 | 0.8522 | 0.8522 | 0.8522 | 0.8522 | 0.8435 | 0.8464 | 0.8522 | 0.8638 |
| | $D6: \frac{1}{k^3}$ | 0.8621 | 0.8609 | 0.8522 | 0.8464 | 0.8464 | 0.8406 | 0.8522 | 0.8609 | 0.8551 |

