## Appendix 6 for "Grand Biological Universe: Genome space geometry unravels looking for a single metric is likely to be futile in evolution": readme.txt

Genome convex hull analysis and tree construction with 24719 sequences in 425 families

Considering the optimal metric in Bacteria

The node name of the tree is "family--class". 

It can be seen that overall the sequences of the same class are relatively close in the tree.
