## Appendix 6 for "Grand Biological Universe: Genome space geometry unravels looking for a single metric is likely to be futile in evolution": readme.txt

Mitochondrial convex hull analysis and tree construction with 253 sequences in 36 families

Considering the optimal metric in fungi: D2, 9-L1

The node name of the tree is "family--class". 

It can be seen that overall the sequences of the same class are relatively close in the tree.

In addition, fungi_data_final(Ver.0515) provides the results of convex hull and 1NN using fungi mitochondrial sequences.
