## Appendix 6 for "Grand Biological Universe: Genome space geometry unravels looking for a single metric is likely to be futile in evolution": readme.txt

Mitochondrial convex hull analysis and tree construction

Considering the optimal metric in invertebrate

The node name of the tree is "family--class".

It can be seen that overall the sequences of the same class are relatively close in the tree.
