## Appendix 6 for "Grand Biological Universe: Genome space geometry unravels looking for a single metric is likely to be futile in evolution": readme.txt

Mitochondrial convex hull analysis and tree construction with 309 sequences in 75 families (some families have no class/order labels)

Considering the optimal metric: L2, 1/k^1.5, 1-9mer averaged natural vectors

The node name of the tree is "family--class". In addition, Phaeophyceae protists (8 families in 3 orders) are analyzed separately and the node name is "family--order".

It can be seen that overall the sequences of the same class/order are relatively close in the tree.
