## Appendix 6 for "Grand Biological Universe: Genome space geometry unravels looking for a single metric is likely to be futile in evolution": readme.txt

Mitochondrial convex hull analysis and tree construction with 5214 sequences in 746 families (as well as a few Ascidian sequences)

Considering the optimal metric: L1, 1/2^k, 1-7mer averaged natural vectors

The node name of the tree is "family--class". In addition, Reptiles (54 families in 4 orders) are analyzed separately and the node name is "family--order".

It can be seen that overall the sequences of the same class/order are relatively close in the tree.
