## Supplementary figures and images for "Grand Biological Universe: Genome space geometry unravels looking for a single metric is likely to be futile in evolution"

### chloroplast_family_ophyta.pdf

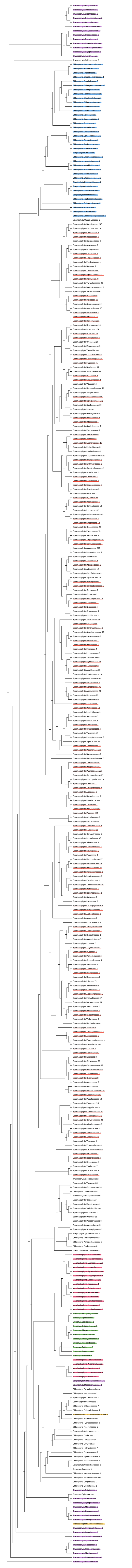
